# Structure-aided development of small molecule inhibitors of ENPP1, the extracellular phosphodiesterase of the immunotransmitter cGAMP

**DOI:** 10.1101/2020.05.30.125534

**Authors:** Jacqueline A Carozza, Jenifer A. Brown, Volker Böhnert, Daniel Fernandez, Yasmeen AlSaif, Rachel E. Mardjuki, Mark Smith, Lingyin Li

**Affiliations:** Department of Chemistry, Stanford University, CA 93405, USA; Department of Stanford ChEM-H, Stanford University, CA 93405, USA; Department of Biophysics Program, Stanford University, CA 93405, USA; Department of Biochemistry, Stanford University, CA 93405, USA; Department of Stanford ChEM-H Macromolecular Structure Knowledge Center, Stanford University, CA 93405, USA; Department of Biology, Stanford University, CA 93405, USA; Department of Stanford ChEM-H Medicinal Chemistry Knowledge Center, Stanford University, CA 93405, USA

## Abstract

Cancer cells initiate an innate immune response by synthesizing and exporting the small molecule immunotransmitter cGAMP, which activates the anti-cancer Stimulator of Interferon Genes (STING) pathway in the host. An extracellular enzyme, ectonucleotide pyrophosphatase phosphodiesterase 1 (ENPP1), hydrolyzes cGAMP and negatively regulates this anti-cancer immune response. Small molecule ENPP1 inhibitors are much needed as tools to study basic biology of extracellular cGAMP and as investigational cancer immunotherapy drugs. Here, we surveyed structure-activity relationships around a series of cell-impermeable and thus extracellular-targeting phosphonate inhibitors of ENPP1. Additionally, we solved the crystal structure of an exemplary phosphonate inhibitor to elucidate the interactions that drive potency. This study yielded several best-in-class compounds with *K*_i_ < 2 nM and excellent physicochemical and pharmacokinetic properties. Finally, we demonstrate that an ENPP1 inhibitor delays tumor growth in a breast cancer mouse model. Together, we have developed ENPP1 inhibitors that are excellent tool compounds and potential therapeutics.

## Introduction

Adaptive immune checkpoint inhibitors such as anti-PD-1, anti-PD-L1, and anti-CTLA-4 are now curing cancer patients who were previously considered terminally ill.^1^ These inhibitors work by removing the immunological brakes that cancer cells place on tumor-infiltrating lymphocytes (TILs), therefore increasing the cancer-killing efficacy of TILs. However, only “hot” tumors – those that already have high numbers of TILs – respond to checkpoint inhibitor therapy. Most tumors do not exhibit this TIL inflammation and thus are immunologically “cold.”^2–4^ Turning “cold” tumors “hot” by activating the innate immune detection of cancer, which is upstream of the recruitment of adaptive immune TILs, could revolutionize cancer immunotherapy.

The cytosolic double-stranded DNA (dsDNA) sensing-Stimulator of Interferon Genes (STING) pathway is the key innate immune pathway that responds to cancerous cells. Chromosomal instability and extrachromosomal DNA are hallmarks of cancer that can lead to leakage of dsDNA into the cytosol.^5–9^ The cytosolic dsDNA is detected by the enzyme cyclic-GMP-AMP synthase (cGAS),^10^ which synthesizes the cyclic dinucleotide 2’,3’-cyclic GMP-AMP (cGAMP).^11,12^ cGAMP then binds and activates STING, which leads to production of Type I interferons (IFNs) and downstream TIL infiltration. We recently discovered that cancer cell lines basally synthesize cGAMP and export it to the extracellular space.^13^ Extracellular cGAMP is then internalized by host cells to trigger anti-cancer innate immune responses.^13–18^ However, we also discovered that the ubiquitously expressed extracellular enzyme ENPP1, which was previously known only as an ATP hydrolase, is the dominant hydrolase for extracellular cGAMP and dampens innate responses to cancer.^13,19^ We previously developed phosphorothioate cGAMP analogs that are resistant to ENPP1 hydrolysis.^19^ Since then, other stable cGAMP analogs have entered clinical trials in combination with anti-PD-1 checkpoint blockers (Trial IDs NCT03172936 and NCT03010176, respectively). However, these cGAMP analogs need to be injected directly into the tumors to achieve efficacy and to avoid systemic interferon response. Alternatively, we propose to inhibit ENPP1 to boost the efficacy of the endogenous extracellular cGAMP that is only locally exported by cancer cells. Indeed, genetic knockout and pharmacological inhibition of ENPP1 both increased tumor-infiltrating dendritic cells and controlled tumor growth, without any safety concerns.^13^ This work suggests that ENPP1 inhibitors could turn “cold” tumors “hot” and render them more sensitive to adaptive immune checkpoint inhibitors.

Outside of its role in cancer immunotherapy, cGAMP has shown excellent adjuvant activities in vaccination against influenza^20^ and may aid in the current urgent demand to develop vaccines against SARS-CoV-2. We, therefore, propose that ENPP1 inhibitors may also stabilize cGAMP when administered as an adjuvant.

ENPP1 is a single-pass transmembrane protein that is anchored with the catalytic domain outside of the cell. It can also be exported extracellularly in a soluble form that is abundant in the circulation.^21–23^ ENPP1 was known to convert nucleotide triphosphates (preferentially ATP) to nucleotide monophosphates (cite something). We reported that ENPP1 also converts cGAMP to AMP and GMP by hydrolyzing first the 2’-5’ phosphodiester bond, and then the 3’-5’.^19^ Similar to the other NPP family members, its catalytic site coordinates two zinc ions that hold the substrate phosphate in place for threonine-mediated nucleophilic attack. The adenosine base of ENPP1 substrates stacks with aromatic residues that form a tight pocket in the active site.^24^ Later co-crystal structures of ENPP1 with pApG, the degradation intermediate of 2’3’-cGAMP, and with 3’3’-cGAMP provided mechanistic insight into why ENPP1 degrades 2’3’-cGAMP, but not 3’3’-cGAMP.^25^ However, all natural ENPP1 substrates have *K*_m_ values higher than 20 μM.^19,24^

Despite ENPP1 being a highly sought-after target,^26–34^ developing drug-like ENPP1 inhibitors has proved difficult. Here, we report the development of the most potent ENPP1 inhibitors to date and the co-crystal structure of an exemplary phosphonate inhibitor with ENPP1. Inspired by the molecular scaffold of a previous inhibitor, QS1,^26,28^ which lacks potency at physiological conditions, we build structure-activity relationships (SAR) around the three sections of the molecule – the zinc-binding head, the core, and the tail – and develop several inhibitors with nanomolar *K*_i_ values. Our crystal structure revealed extensive interactions between the inhibitor and ENPP1 and explains its 1000-10,000-fold improvement in affinity over the natural substrates. Finally, these inhibitors have desirable physicochemical and pharmacokinetic properties, which enables their systemic use in mouse studies. Indeed, we demonstrate that treatment with one of our top ENPP1 inhibitors delays tumor growth in a breast cancer mouse model.

## Results

### cGAMP is rapidly degraded in human and mouse plasma

Given that a soluble form of ENPP1 is present in the circulation, we first asked how much ENPP1 activity is present in freshly drawn mouse and human plasma by measuring the half-life of radiolabeled cGAMP. cGAMP was degraded rapidly (*t*_1/2_ = 16 minutes for mouse; *t*_1/2_ = 30–60 minutes for human, based on 5 healthy donors) (Fig 1a–c). Detectable hydrolysis of cGAMP ex vivo occurs only in WT, and not *Enpp1*^-/-^ mouse plasma^19^, and inhibiting ENPP1 activity using EDTA to chelate the catalytic zinc ions abrogated cGAMP degradation in both mouse and human plasma (Fig 1a–c). Rapid degradation of extracellular cGAMP by circulating ENPP1 underscores the need to develop potent and systemic ENPP1 inhibitors.

**Figure 1.**
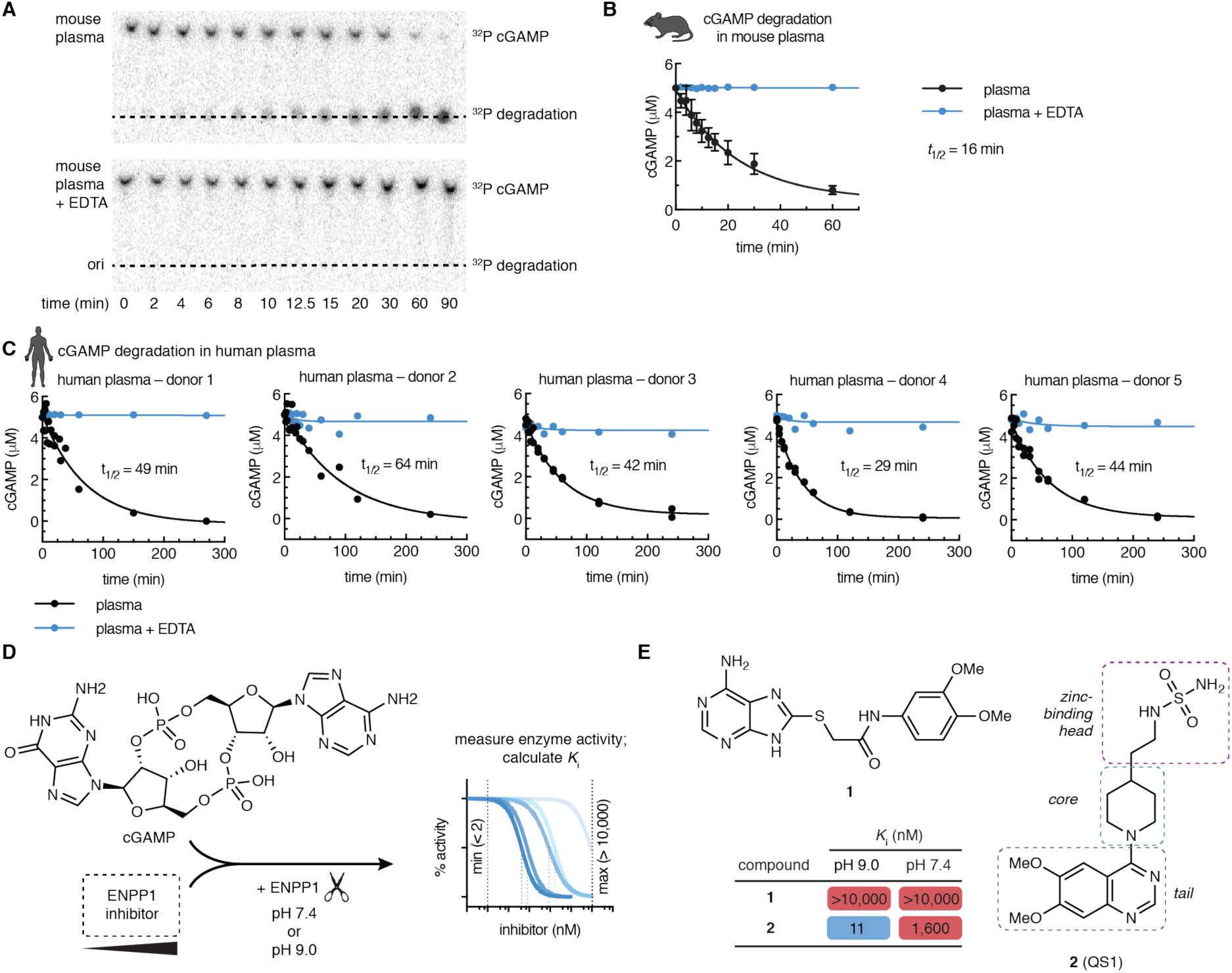
ENPP1 rapidly degrades cGAMP but current inhibitors are not potent enough as tools to investigate ENPP1 function. (A)–(C) cGAMP (5 μM, with trace radiolabeled [^32^P]cGAMP) was incubated with C57B6/J mouse plasma (A, B) or human plasma (C) at 37 °C. 50 mM EDTA was included in the indicated reactions. At indicated times, degradation was assessed by separation of cGAMP from degradation products by thin layer chromatography (TLC) and autoradiography. (A) shows a representative experiment for mouse plasma. For (B), data are from 4 independent experiments, mean + SD shown. For (C), data are from 3 donors, each with 2 independent experiments; replicates are plotted individually. (D) Assay for measuring ENPP1 inhibitor *K*_i_ values. cGAMP (5μM) was incubated with increasing concentrations of ENPP1 inhibitor (maximum = 10 μM) and ENPP1 (3 nM) in a buffer containing 50 mM Tris, 150 mM NaCl, 1 μM ZnCl_2_, and 500 μM CaCl_2_ at the indicated pH value. The minimum measurable *K*_i_ is 2 nM (determined by enzyme concentration); the maximum is 10 μM. (E) Chemical structures and Ki values of compounds **1** and **2** (mean of at least 2 independent replicates; evaluated in assay described in D).

### Previous ENPP1 inhibitors are not potent enough as tools to investigate ENPP1 function

Assays used to assess the potency of previously attempted ENPP1 inhibitors are inconsistent^26–34^; however, developing an appropriate assay is key to determining the utility of the molecules in inhibiting cGAMP degradation under physiological conditions. The most common model substrate is *p*-nitrophenyl-5’-TMP (*p*-NPTMP), but IC_50_ values measured using this substrate have been shown to deviate significantly from IC_50_ values measured using a natural substrate such as ATP.^30^ In addition, perhaps to increase the speed of the assay, most reported assays are conducted at pH 9, where ENPP1 is most active. However, since ENPP1 is active in serum at physiological pH (Fig. 1a–c), an effective ENPP1 inhibitor needs to be potent at pH 7.4 or even lower, as can occur in the tumor microenvironment.^35,36^ Therefore, we evaluated inhibitors with an assay using cGAMP as a substrate at pH 7.4 (Fig. 1d).^37^

There were only two non-nucleotide inhibitor hits from the literature which had the potential to be developed into a lead compound for inhibiting ENPP1. First, a thioacetamide inhibitor (compound **1**, Fig. 1e) was reported to have a *K*_i_ of 5 nM against human ENPP1 using *p*-nitrophenyl-5’-TMP (*p*-NPTMP) as the substrate, but the *K*_i_ increased to 5 µM when using ATP as the substrate, both assays conducted at pH 9.^27^ When we tested the potency of compound **1** using mouse ENPP1 and cGAMP as a substrate at both pH 7.4 and pH 9, we detected no activity at concentrations below 10 µM (Fig. 1e), suggesting that it cannot block cGAMP degradation activity specifically compared to other substrates, or that it may be specific to human ENPP1. Regardless, lack of efficacy against cGAMP and/or mouse ENPP1 disqualifies this molecule as a scaffold to use for further development.

Second, Patel et al reported a quinazoline-piperidine-sulfamide inhibitor (QS1, compound **2**) with an IC_50_ of 36 nM against ENPP1 using ATP as a substrate.^26^ However, we found that the potency of QS1 dropped 100-fold when adjusted the pH down to 7.4 instead of pH 9 (*K*_i_ = 1.6 μM using cGAMP as a substrate, Fig. 1e), making it unsuitable for experiments at physiological pH and in vivo. In addition, we have previously shown that QS1 non-specifically blocks cGAMP export, limiting its use as a tool for studying extracellular cGAMP biology.^13^ We, therefore, sought to develop more potent and specific ENPP1 inhibitors.

### ENPP1 inhibitors with a zinc-binding phosphonate head group exhibit high potency at physiological pH

We hypothesized that QS1 lacked potency at pH 7.4 because deprotonation of the sulfamide group (*p*K_a_ around 7–8) is important for efficient binding to the zinc atoms at the catalytic site of ENPP1. Therefore, we tested several other classes of zinc-binding head groups on the QS1 scaffold (Fig. 2a–b). We also tried methylene, ethyl, and/or propyl linkage between the head groups and the piperidine core. Ureas (compounds **4** and **5**) and carboxylic acid (compound **6**) were less potent compared to sulfamides (compounds **2** and **3**), but we observed improvement in potency with boronic acid (compound **7**), hydroxamic acids (compounds **8** and **9**), and especially phosphates (compounds **10** and **11**), the natural zinc binding group of the ENPP1 substrates cGAMP and ATP. To increase compound stability while preserving the core properties of phosphates, we tested thiophosphates (compounds **12** and **13**), phosphonates (compounds **14**–**16**), and a thiophosphonate (compound **17**), all of which are known to be more stable against enzymatic degradation compared to phosphates. The phosphonates showed *K*_i_ values less than 50 nM independent of the pH (Fig. 2a–b).

**Figure 2.**
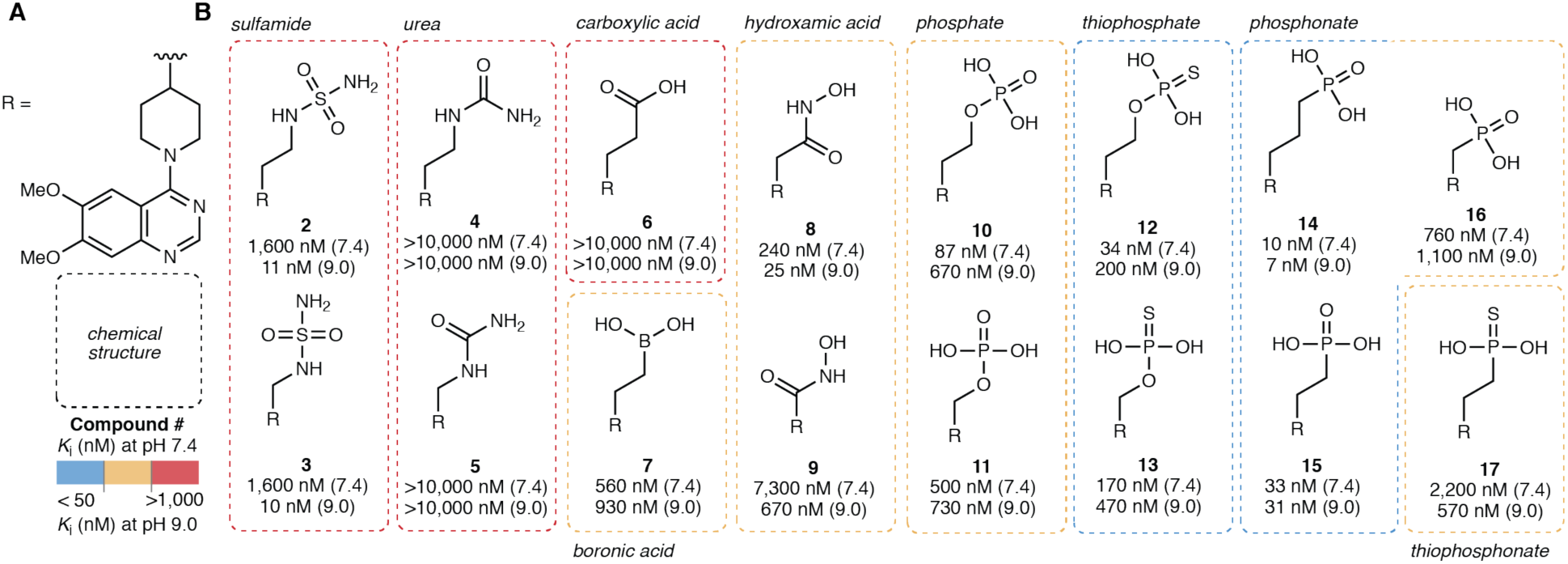
ENPP1 inhibitors with a zinc-binding phosphonate head group exhibit high potency at physiological pH. (A) Chemical structure of the R group (core = piperidine, tail = 6,7-dimethoxy quinazoline). (B) Chemical structures of zinc-binding heads with corresponding *K*_i_ values (mean of at least 2 independent replicates) at pH 7.4 and pH 9.0. *K*_i_ values were determined using 3 nM ENPP1 and 5 μM cGAMP.

We chose to proceed with phosphonates because they are potent, stable, and synthetically tractable. In addition, there are several phosphonate drugs on the market including Tenofovir and Pradefovir as antivirals, Fosfomycin as an antibiotic, and bisphosphonates (Pamidronic aicd, Zoledronic acid, Alendronic acid) for osteoporosis and bone disease. Finally, they are negatively charged at all physiological pH values, which is crucial for zinc binding and keeping them cell impermeable to act on the extracellular target ENPP1. We also chose the ethylene linker (compound **15**, *K*_i_ = 33 nM, previously reported as STF-1084^13^) because it has been shown previously that for the sulfamides, analogs with shorter linker lengths have less affinity for the cardiac potassium channel hERG, a detrimental off target.^28,32^

### Co-crystal structure of ENPP1 and compound 15 reveals molecular determinants of potency

ENPP1 is a >100 kDa multidomain glycoprotein with three glycosylation chains. It has a catalytic domain, a nuclease-like domain which provides structural support important for catalysis, and two disordered SMB domains. We used a mouse ENPP1 construct where the transmembrane anchor was truncated and replaced by a signal peptide, and expressed the construct in HEK293S GnT1^-^ cells to generate secreted soluble ENPP1 with simplified glycosylation chains as previously described^24,25,38^. We then determined the structure of ENPP1 in complex with compound **15** to 3.2 Å resolution using X-ray crystallography (Fig. 3, Fig. S1, Tables S1–3). Compound **15** occupies the substrate-binding pocket (Fig. 3a–d) and forms extensive interactions with zinc ions and residues in the catalytic site, demonstrating that compound **15** is a competitive inhibitor. The phosphonate oxygens bind both zinc ions, and the third phosphonate oxygen forms a hydrogen bond with N259, a residue previously determined to be important for catalytic activity. The piperidine group is engaged in hydrophobic interactions with L272, and the quinazoline group sits between Y322 and F239 to form π-π interactions with both (Fig. 3b–d). Although compound **15** adopts a similar binding conformation as the reaction product AMP^24^ (PDB 4GTW, Fig. 3e–f), there are two significant differences between the ligands that suggest why compound **15** has a much higher affinity than the substrates ATP or cGAMP. First, the quinazoline ring of compound **15** sits ∼2.5 A further back in the catalytic pocket compared to the purine ring of AMP, perhaps facilitating hydrophobic interactions with residues in the back of the pocket, and possibly a polar interaction between N3 of the quinazoline and Y353. Second, the 7-methoxy group of compound **15** makes a direct hydrogen bond with D308, whereas AMP makes a water-mediated hydrogen bond (Fig. 3e). In addition, residue K277 forms a hydrogen bond with the 7-methoxy oxygen, further strengthening this direct hydrogen bond network. Together, the potency of compound **15** is driven by its zinc ion binding, hydrophobic interactions with the back of the binding pocket, and direct hydrogen bonds with active side resides of ENPP1. In addition, this structure guided our SAR for the core and tail parts of the inhibitor.

**Figure 3.**
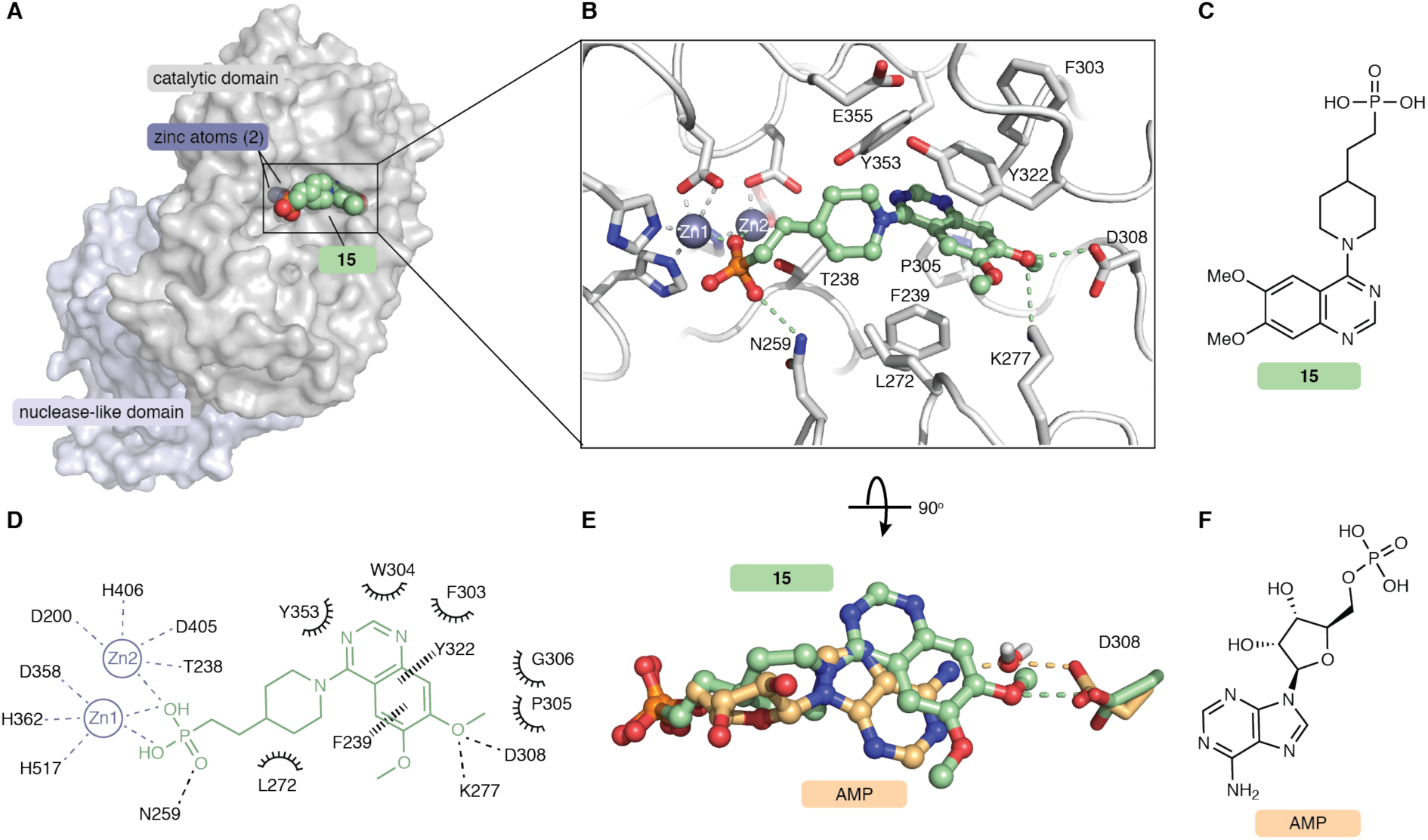
Co-crystal structure of ENPP1 and compound 15 reveals molecular determinants of potency. (A) The 3.2 Å crystal structure of compound **15** (green spheres) bound to mouse ENPP1 (gray surface; catalytic and nuclease-like domains). (B) Expanded view of compound **15** (green sticks/spheres) bound in the active site of ENPP1 (gray sticks). Zincs are shown as dark gray spheres. Hydrogen bonds and metal coordination are shown as dashed lines. (C) Chemical structure of compound **15**. (D) Schematic drawing of interactions formed between compound **15** (green) and the ENPP1 active site (black). Residues within 5 Å are shown. Metal coordination shown as gray dashed lines, hydrogen bonds shown as black dashed lines, aromatic interactions shown as black wedged lines, and hydrophobic or polar interactions shown as spokes. (E) Overlay of compound **15** (green) with the AMP (orange, PDB 4GTW), both bound to ENPP1. Ligands are shown as sticks/spheres, protein residue D308 is shown as sticks, and water is shown as sticks/spheres. Density for water molecule is present only in AMP-ENPP1 crystal structure. (F) Chemical structure of AMP.

### Piperidine and benzyl amine cores are optimal molecular linkers

Since our crystal structure suggests that the zinc-binding phosphonate head and quinazoline tail form the most important interactions with ENPP1, we next sought to explore the core region to achieve optimal geometry between these two functional groups (Fig. 4a–b). Inverting the nitrogen in the piperidine from aryl to alkyl (compound **18**) resulted in more than a 200-fold loss in potency, as did installing a piperazine (compound **19**). The benzyl group in compound **20** resulted in approximately a 3-fold loss in potency compared to the piperidine in compound **15**. When we installed a nitrogen to form aminobenzyls (compounds **21** and **22**), we again observed small losses in potency. Extending the length to a benzyl amine (compounds **23** and **24**) improved *K*_i_ values to 12 nM and 26 nM, respectively. We anticipate that the core region itself does not make significant contacts with ENPP1, but rather dictates the angle between head and tail. The piperidine (compound **15**) and benzyl amine (compounds **23** and **24**) cores have unique geometry but both result in low nanomolar *K*_i_ values, suggesting that these linkers yield favorable binding poses.

**Figure 4.**
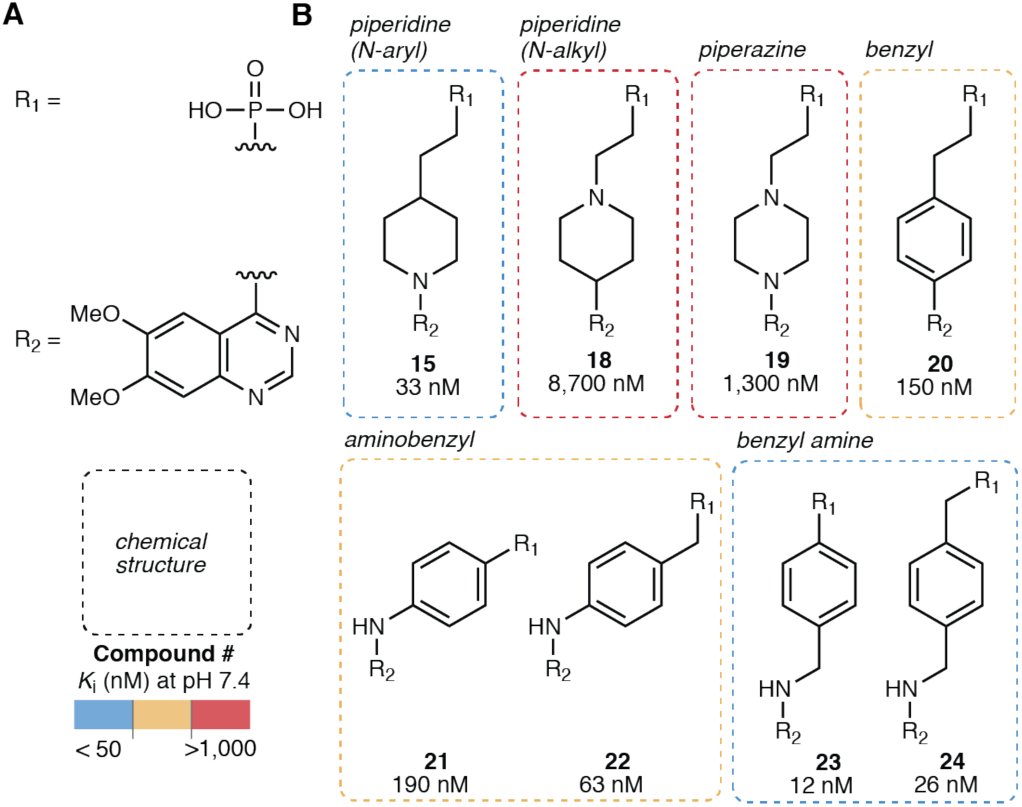
Piperidine and benzyl amine cores are optimal molecular linkers. (A) Chemical structures of the R_1_ group (phosphonate head) and the R_2_ group (6,7-dimethoxy quinazoline tail). (B) Chemical structures of cores with corresponding *K*_i_ values (mean of at least 2 independent replicates) at pH 7.4. *K*_i_ values were determined using 3 nM ENPP1 and 5 μM cGAMP.

### 8-methoxy quinazoline tails achieve high potency compared to other quinazoline substitutions

Using the phosphonate zinc-binding head and the piperidine core as a scaffold, we then sought to determine optimal substitution of the quinazoline tail (Fig. 5a–b). Based on our co-crystal structure of ENPP1 with compound **15**, we predicted that the 7-methoxy would be critical for binding to ENPP1, while the 6-methoxy would be dispensable since it is solvent-exposed. Indeed, when we deleted the methoxy groups individually, we found that the 6-methoxy alone (compound **25**) was 100-fold less potent, while the 7-methoxy alone (compound **27**) displayed similar *K*_i_ to the 6,7-methoxy (compound **15**). Substituting a hydrogen bond donor/electron-donating group in the 7 position (compounds **29** and **30**) led to a loss of potency compared to the 7-methoxy (compound **27**), suggesting that the methyl group makes favorable interactions with the protein. Extending to a 7-ethoxy (compound **28**) or 7-isopropyloxy (compound **31**) was not tolerated, presumably due to steric constraints.

**Figure 5.**
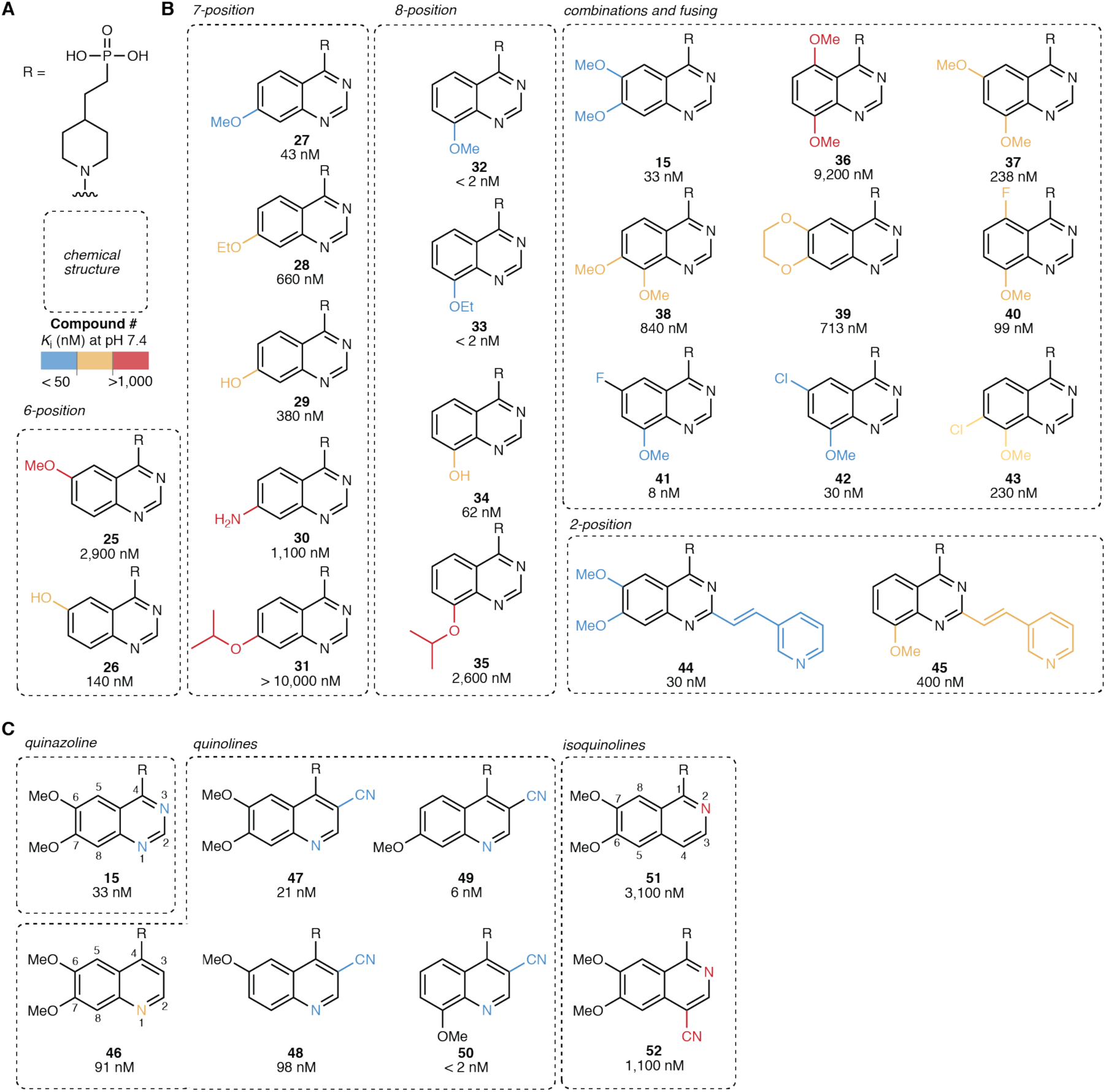
8-methoxy quinazoline and 3-nitrile quinoline tails achieve high potency. (A) Chemical structures of the R group (head = phosphonate, core = piperidine). (B)–(C) Chemical structures of quinazoline (B) and quinoline (C) tails with corresponding *K*_i_ values (mean of at least 2 independent replicates) at pH 7.4. *K*_i_ values were determined using 3 nM ENPP1 and 5 μM cGAMP.

We then moved to substituents at the 8-position of the quinazoline ring. The 8-methoxy (compound **32**) and 8-ethoxy (compound **33**) quinazolines displayed impressive potency with a *K*_i_ values less than 2 nM, which is the limit of quantification of the assay when using a 3 nM concentration of ENPP1. We reasoned that combining other methoxy groups with the 8-methoxy could make additional interactions with the binding pocket, since the 7-methoxy (compound **27**) alone was also potent. However, combinations of 8-methoxy with substituents in either the 5, 6, or 7 positions (compounds **36**–**38** and **40**–**43**) all decreased potency. From these data and the crystal structure, we hypothesize that the 8-methoxy (compound **32**) and 8-ethoxy (compound **33**) quinazolines make hydrogen bonds with both D208 and/or K277 similar to the 7-methoxy in compound **15**. This would suggest that compounds **32** and **33** shift in the pocket to accommodate these interactions. Adding more substituents to the quinazoline does not necessarily lower potency further, as the inhibitor-interacting residues may already be occupied.

In addition, similar to previous SAR,^26,32^ the 2-substituted vinyl-3-pyridyl combined with the 6,7-dimethoxy (compound **44**) showed identical potency to its parent 6,7-dimethoxy (compound **15**). This is surprising, since the crystal structure shows limited space in this area of the pocket, suggesting that the pyridine may make a specific interaction with a protein residue. However, this trend did not hold for the 8-methoxy (compound **45**), where the addition of the 2-substituted vinyl-3-pyridyl decreased potency by more than 100-fold. This further supports the hypothesis that different substitutions cause shifts in the inhibitor binding to the pocket, possibly excluding space that was previously available.

### 3-nitrile quinolines substitute for quinazolines in the tail

From the crystal structure of compound **15** bound to ENPP1, we noted that the quinazoline tail of compound **15** is close to residues near the back of the binding pocket. We perturbed the tail aromatic ring structure itself to investigate these possible interactions (Fig. 5a,c). Working from the quinazoline compound **15**, we deleted the nitrogens on the ring on by one. Deleting N3 (quinoline, compound **46**) led to merely a 3-fold decrease in potency, but deleting N1 (isoquinoline, compound **51**) led to a more than 100-fold decrease in potency. We tested whether we could replace the inhibitor-protein interaction that involved the N3 or N1 with a nitrile, a more electronegative group and possible hydrogen bond acceptor, as has been previously suggested.^39^ Indeed, the 6,7-dimethoxy, 3-nitrile quinoline (compound **47**) was more potent than the quinazoline itself (compound **46**), and this pattern repeated with all of the other methoxy substituents (compounds **48**–**50**). Compound **50** was the most potent molecule with a *K*_i_ less than our limit of quantification of 2 nM. When we attempted to substitute back the missing nitrogen with a nitrile group on the isoquinoline scaffold (compound **52**), we lost potency, possibly due to molecular clashes between inhibitor and protein. In summary, the N1 nitrogen is critical for potency, possibly due to polar interactions with the protein backbone ∼4 Å away. It cannot be replaced by a nitrile since space is limited. The N3 nitrogen can be replaced by a more electronegative nitrile, leading to much more potent compounds.

### Hybrid ENPP1 inhibitors combine the most potent heads, cores, and tails

Taking into account of all of our SAR data, we then synthesized hybrid molecules composed of the most potent heads, cores, and tails (8-methoxy quinazoline, e.g., compound **32**, and 8-methoxy quinoline 3-nitrile, e.g., compound **50**) (Fig. 6). For all of these molecules, *K*_i_ values fell below 100 nM. Combining the benzyl amine core with the 8-methoxy quinoline 3-nitrile tail (compound **57**) yielded another inhibitor with a *K*_i_ less than our limit of quantification of 2 nM. Attaching other zinc binding head groups, including thiophosphate (compound **53**), boronic acid (compounds **60** and **61**), and hydroxamic acid (compound **62**), to the most potent core/tail combinations also yielded potent inhibitors.

**Figure 6.**
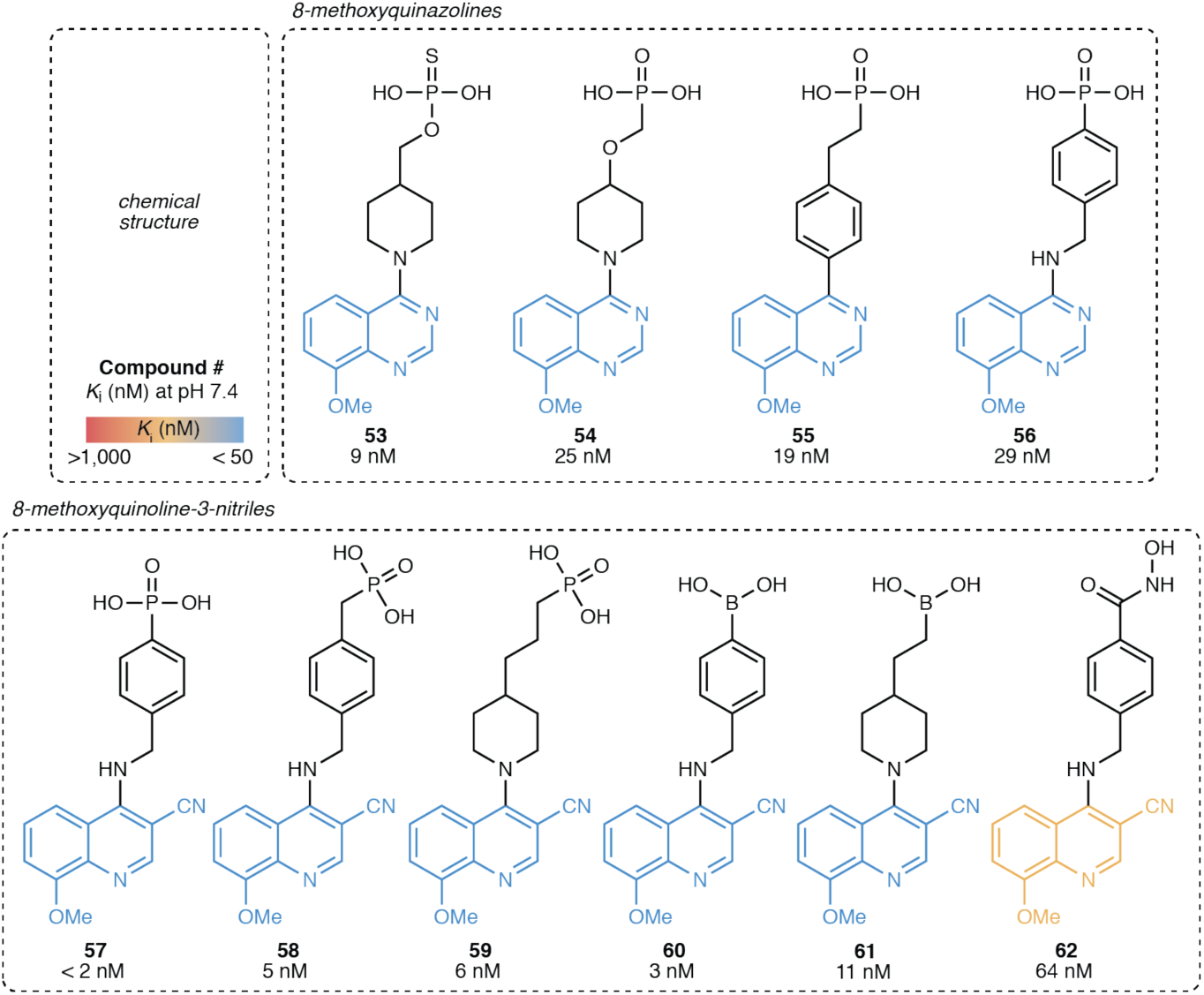
Hybrid ENPP1 inhibitors combine the most potent heads, cores, and tails. Chemical structures of hybrid ENPP1 inhibitors connecting the most potent heads, cores, and tails with corresponding *K*_i_ values (mean of at least 2 independent replicates) at pH 7.4. *K*_i_ values were determined using 3 nM ENPP1 and 5 μM cGAMP.

### Compound 32 leads to tumor growth delay in mouse breast cancer model after optimizing in vitro ADME and in vivo pharmacokinetics

We further evaluated the potency and in vitro ADME properties of the seven most potent inhibitors. First, we evaluated the protein-shifted IC_50_, which can help predict what the efficacy will be in vivo, as this value is generally higher than that observed in vitro due to protein binding. We observed that all of the inhibitors displayed a shift in potency when we did the assay in the presence of human serum albumin, although several still stayed below 50 nM (Fig. 7a, Table 1, Fig. S2). We then tested their potency in both mouse and human plasma, and obtained similar values, confirming that mid-nanomolar concentrations are sufficient in biological fluids to prevent degradation of cGAMP (Fig. 7b, Table 1). Although ENPP1 protein is present in our circulation, these data demonstrate that we can completely block serum ENPP1 activity with as little as 100 nM of our most potent inhibitors, showing that protein binding does not negate the inhibition, and suggesting that serum ENPP1 will not sequester all the systemically administered ENPP1 inhibitor. Finally, all of our inhibitors are stable and cell impermeable. Although we expect these phosphonate inhibitors to have few intracellular off-targets due to their impermeability, we also confirmed that the inhibitors are non-toxic to primary human peripheral blood mononuclear cells (PBMCs) (Table 1, Fig. S3). Since QS1 has an off-target hERG liability,^26^ we tested two phosphonate inhibitors (compounds **15** and **32**) and observed no inhibition at 25 μM (Table 1).

**Table 1:** Potency and in vitro ADME characterization of top ENPP1 inhibitors.

| Assay | Compound |  |  |  |  |  |  |
| --- | --- | --- | --- | --- | --- | --- | --- |
|  | 15 | 32 | 33 | 49 | 50 | 57 | 59 |
| $K_i$ (nM) | 33 | < 2 | < 2 | 6 | < 2 | < 2 | 6 |
| $K_i$ (HSA, nM) | 163 | 6 | 4 | 182 | 6 | 5 | 31 |
| IC<sub>50</sub> mouse plasma (nM) | 97 | 4 | 8 | n.d. | n.d. | 33 | 36 |
| IC<sub>50</sub> human plasma (nM; donor 1/2/3) | 25/66/54 | 6/4/8 | 10/4/7 | n.d. | n.d. | 18/11/18 | 29/140/70 |
| CC<sub>50</sub> (PBMCs, μM) | > 100<sup>13</sup> | > 100<sup>13</sup> | > 100 | > 100 | > 100 | > 100 | > 100 |
| Microsome stability (t<sub>1/2</sub>) (min; h/m) | >159 / >159<sup>13</sup> | >159 / >159<sup>13</sup> | >159 / >159 | >159 / >159 | >159 / >159 | >159 / 147 | >159 / >159 |
| Caco-2 (1 $\times 10^{-6}$ cm/s) | 0.47<sup>13</sup> | 0.1<sup>13</sup> | 0.2 | 0.0 | 0.1 | 0.2 | 0.5 |
| Protein Binding (%; h/m) | 71.2/31.2 | 66.2/55.0 | 62.0/47.0 | 93.7/48.3 | 78.1/26.9 | 89.5/27.9 | 89.6/51.6 |
| IC<sub>50</sub> hERG (μM) | >25 | >25 | n.d. | n.d. | n.d. | n.d. | n.d. |

**Figure 7.**
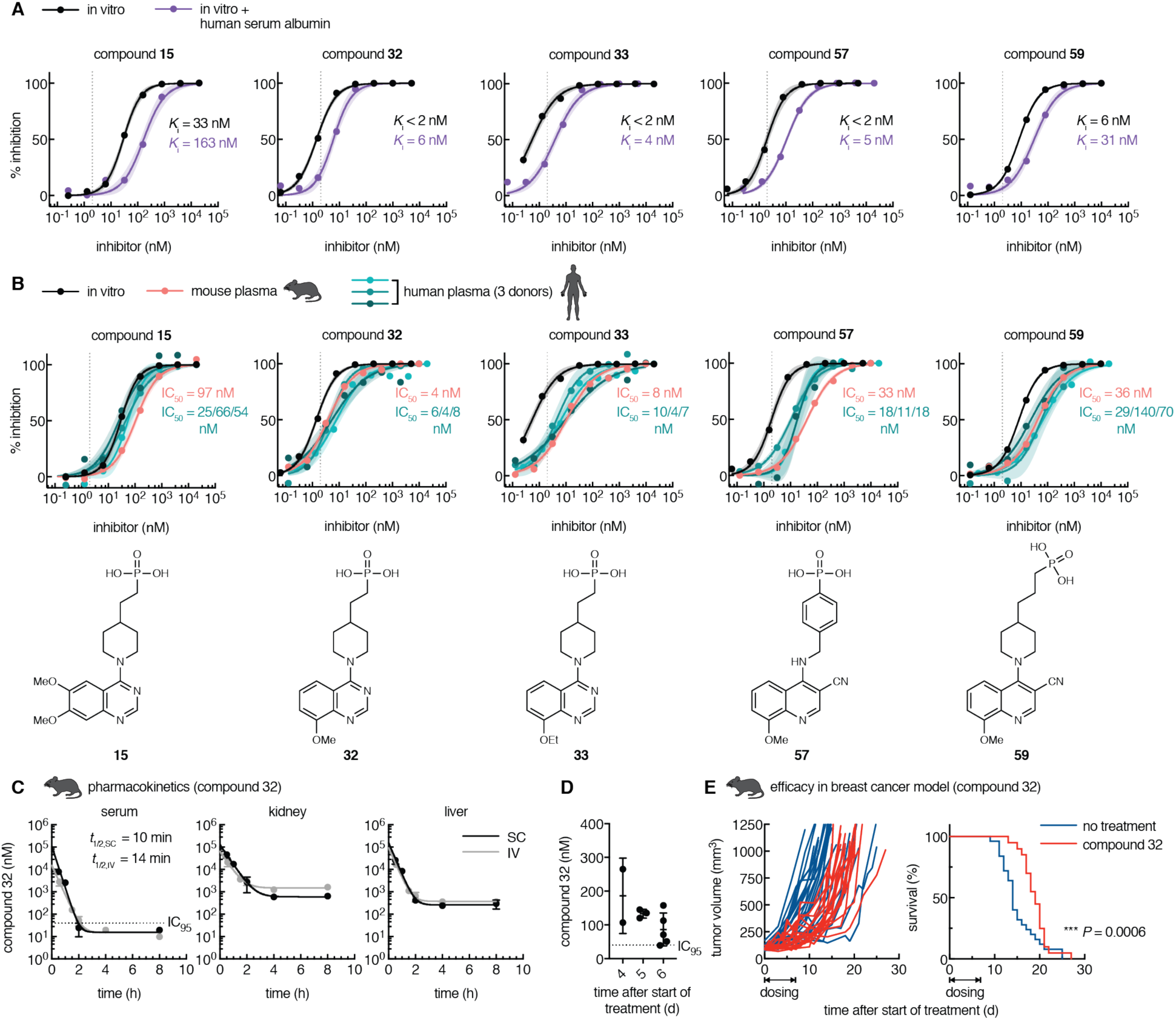
Optimized phosphonate ENPP1 inhibitors prevent cGAMP degradation in mouse and human plasma at nanomolar concentrations. (A) In vitro dose-inhibition curves for selected top ENPP1 inhibitors. *K*_i_ values were determined using 3 nM ENPP1 and 5 μM cGAMP and (where indicated) in the presence of 40 μM human serum albumin at pH 7.4. Dotted line represents the minimum *K*_i_ value (2 nM) measurable with the given enzyme concentration. (B) In mouse and human plasma, IC_50_ values were determined using 5 μM cGAMP with trace radiolabeled [^32^P]cGAMP. In vitro dose-response curves replotted from (A). Chemical structures of inhibitors are displayed below. Dots represent the mean of two independent replicates, and shaded areas around the fitted curves represent the 95% confidence interval of the fit. (C) Concentrations of compound 32 in mouse serum, kidney, and liver after one 10 mg/kg dose via intravenous (IV) or subcutaneous (SC) route. IC_95_ = 40 nM; LOQ = 10 nM. Mean ± SEM is plotted, *n* = 2 for each point except the following, where *n* = 1: liver SC 4 h (point removed as an outlier using the ROUT method with Q = 1%) and serum 8 h (concentration below LOQ). Data were fit with one-phase decay with 1/Y^2^ weighting to obtain half-life. (D) Concentrations of compound 32 in mouse serum after consecutive once daily dosing at 300 mg/kg via SC route for 6 days. Serum was collected for analysis 24 hours after last dose, immediately before the next dose. (E) Tumor volumes (left; each line represents one mouse) and survival (right) of mice bearing E0771 breast cancer tumors. Mice were treated with vehicle (no treatment, *n* = 25) or treated with compound 32 (*n* = 20) dosed once daily at 300 mg/kg via SC route for seven consecutive days after tumors reached 80-120 mm^3^. For survival, *P* = 0.0006 (Gehan-Breslow-Wilcoxon test).

We then assessed the pharmacokinetic (PK) profile of one of the top inhibitors, compound 32, in mice. Since compound 32 has an IC_50_ value of 4 nM in mouse plasma, we aim to achieve serum concentrations of 40 nM (IC_95_ value). First, we administered intravenous (IV) and subcutaneous (SC) doses of compound 32 at 10 mg/kg to mice and analyzed the concentration of compound 32 in the serum, kidney, and liver for the next 8 hours. We found that serum concentrations declined to 10 nM or less within 8 hours for both IV and SC administration, and the half-life was only 10-15 minutes. Concurrent with the rapid decline in serum concentrations, we saw concentration plateaus in the kidney and liver in the micromolar range, suggesting that compound 32 is rapidly excreted through these organs. This is in agreement with our previously reported PK study of compound 32 performed at 300 mg/kg via SC administration where we observed that the serum concentration drops to ∼100 nM at the end of 24 hours.^13^ Therefore, we were only able to test the efficacy of compound 32 in that study dosing via an osmotic pump to maintain serum concentration of compound 32 above its IC_95_. To achieve sustained serum concentrations without surgically implanting osmotic pumps, we repeated the 300 mg/kg SC dosing daily for 6 days. We collected serum 24 hours after the previous dose, which corresponds to the expected lowest point in the trough. We observed on average ∼100 nM of compound 32 across several days, and the lowest concentration measured was 40 nM. We also observed no adverse effects and the mice maintained healthy weights, suggesting that this dosage amount and timing is tolerated well. Together, we succeeded in achieving convenient, systemic, once daily dosing of compound 32 that results in sustained serum concentrations above the IC_95_.

After optimization of compound 32 PK, we then asked if compound 32 has anti-tumor efficacy in the orthotopic and syngeneic E0771 triple negative breast cancer model. We chose this model because E0771 cells basally export cGAMP in cell culture. In addition, we previously saw that E0771 cells implanted into mice grow more slowly in *Enpp1*^-/-^ mice than in wild type mice.^13^ We treated mice with established E0771 tumors (∼100 mm^3^) with compound 32 using the optimized dosing schedule for seven consecutive days. Remarkably, we observed a delay in tumor growth in mice treated with compound 32 relative to the controls, which also led to prolonged survival. The tumor growth delay observed with pharmacological inhibition of ENPP1 mirrors that from genetic ablation of ENPP1 we previously reported,^13^ demonstrating that our ENPP1 inhibitors are therapeutically beneficial.

## Discussion

Here we report the development of highly potent phosphonate ENPP1 inhibitors. We demonstrate that they are active against the physiological substrate cGAMP under physiological conditions, including an in vitro assay at pH 7.4 and an ex vivo assay against ENPP1 in mouse and human plasma. Our compounds are by far the most potent inhibitors reported to date. Although some of the boronic acids (compounds **60** and **61**) and hydroxamic acids (compound **62**) are also potent, further investigation of their cell permeability and possible off-target effects would be needed. The lead phosphonate compounds, however, are cell impermeable through passive diffusion, avoiding all potential intracellular off-targets. We, therefore, nominate them as specific tool compounds to study ENPP1 and extracellular cGAMP biology.

To understand the potency of our compounds, we solved the crystal structure of ENPP1 with compound **15**. We found that compound **15** adopts a similar binding pose as AMP, the product of cGAMP and ATP hydrolysis, but there are extra interactions that explain its enhanced potency including a direct hydrogen bond between the 7-methoxy and D308 (instead of water-mediated), extra hydrogen bond interactions with K277, and other hydrophobic and polar interactions with L272 and Y353. Our SAR of other phosphonate inhibitors allowed us to build a model of the drivers of potency. The π-π stacking interactions are key, as only a couple of the cores we tried resulted in potent inhibitors and the core region could be important for positioning of the zinc-binding phosphonate with respect to the π-π stacking tail. We hypothesize that moving the methoxy groups to different positions on the ring would also engage D308 and/or K277 in hydrogen bonding. Even though some of the positions on the ring are solvent-exposed in the compound **15** structure in complex with ENPP1, combinations of more substituents are detrimental to potency, suggesting that these compounds could be shifted in the binding pocket leading to steric hinderance when extra substituents are added. It is interesting to note that ENPP1 makes better interactions with the methoxy group, in contrast to the hydroxy or amine (e.g., compare compound **27** to compounds **29** and **30**; compare compound **32** to compound **34**), suggesting that aliphatic carbons are important in addition to hydrogen bonding. In addition, we can speculate about the importance of the 3-nitrile group off the quinoline, e.g. compound **47** and related analogs. Compared to the N3 on the quinazoline, it is possible that the nitrile can make stronger polar interactions with the protein backbone or surrounding residues (e.g., D200, Y353, or E355), act as a hydrogen bond acceptor, or polarize the quinoline ring for better π-π stacking.^40^ Previous modeling showed that nitriles can replace water-mediated azomethine-protein interactions^39^. This could extend to our scenario in replacing a weak polar interaction with a stronger one. In addition, it is surprising that the 2-vinyl-3-pyridyl substituent (compound **44**) is potent, given that the 2 position faces the protein backbone. It is possible that this large substituent can slide into the narrow pocket between H242 and Y353 or could make a specific interaction with one of the residues. It also suggests that this space is not as available with the 8-methoxy substituent, as compound **45** (with 2-vinyl-3-pyridyl) was more than one hundred-fold less potent than compound **32** (without 2-vinyl-3-pyridyl).

In addition to unmatched potency, our ENPP1 inhibitors also have desirable ADME profiles in vitro, and we have measured the pharmacokinetics of one of the top inhibitors to demonstrate efficacious concentrations in mice with once daily systemic dosing. Compared to jump starting the anti-cancer innate immune response by treating with direct STING agonists, our ENPP1 inhibitors provide two advantages. First, although endogenous extracellular cGAMP enhances the anti-tumor immune response,^13^ the same may not be true for cGAMP analogs. Because cGAMP is specifically transported into different cell types through different transporters,^14,15^ it is difficult to design cGAMP analogs that match the cell-targeting profile of cGAMP itself. It is important to target specific cells because STING activation in cancer cells promotes metastasis,^8^ and STING activation in T cells leads to T cell death,^41–43^ both of which would be detrimental to cancer patients. In contrast, ENPP1 inhibitors should increase the half-life of endogenous cGAMP and thus enhance the natural anti-tumor response. Indeed, treatment of mice with compound 32 delayed tumor growth in the E0771 breast cancer mouse model, which is a promising result for ENPP1 inhibitors as cancer therapeutics. Second, STING agonists can only be introduced intratumorally to achieve efficacy and to avoid systemic inflammation, which limits treatment to palpable and injectable tumors. Since our ENPP1 inhibitors can be administered systemically, we hypothesize that ENPP1 inhibitors could treat a wide variety of cancers and may even be effective against undetectable micrometastasis. They could also avoid causing widespread toxic interferon production, since they only enhance endogenous cGAMP. This hypothesis is supported by the fact that ENPP1-inactivating mutations in both humans and mice do not cause interferonopathy. Our ENPP1 inhibitors are biological tools as well as candidate investigational drugs that have shown efficacy in mouse tumor models, and they hold the promise to potentiate the efficacy of radiation and other immune checkpoint inhibitors. Finally, since cGAMP has shown stunning results as an adjuvant for influenza vaccination,^20^ our lead ENPP1 inhibitor that prevents its extracellular degradation may have the potential to maximize cGAMP’s adjuvancy effect in developing vaccines for pandemic threats.

## Acknowledgments

This work was performed in collaboration with the ChEM-H Medicinal Chemistry Knowledge Center. We thank Dr. Sam Banister, Stephen R. Stabler, Dr. Phil Thomson, and members of the ACME Biosciences synthetic chemistry team for compound synthesis; Khanh Nguyen and Grace Lam for in vitro ADME assays and LC-MS/MS analysis; beamline staff at the Advanced Light Source for technical support during diffraction data collection; and Dr. Dan Herschlag and all members of the Li lab and Medicinal Chemistry Knowledge Center for helpful discussions. This research was supported by the Stanford ChEM-H Medicinal Chemistry Knowledge Center and Macromolecular Structure Knowledge Center and the National Institutes of Health award DP2CA228044 (L.L.), DOD grant W81XWH-18-1-0041 (L.L.), Ono Pharma Foundation Grant (L.L.), NIH fellowship F31CA239510 (J.A.C.), Xu Family Foundation Stanford Interdisciplinary Graduate Fellowship affiliated with ChEM-H (J.A.C.), and National Science Foundation graduate fellowship DGE-1656518 (J.A.B. and R.E.M.). This research used resources of the Advanced Light Source, which is a DOE Office of Science User Facility under contract no. DE-AC02-05CH11231.

## Author contributions

J.A.C., M.S., and L.L. designed the study. J.A.C., Y.A.S., and R.E.M. performed enzyme assays and analyzed the data. V.B. performed mouse experiments. J.A.B. and D.F. determined the crystal structure. J.A.C., M.S., and L.L. wrote the paper. All authors discussed the findings and commented on the manuscript.

## Methods

### Plasma kinetics and IC_50_ value determination

All procedures to obtain plasma after blood draw were performed at 4 °C. Blood from C57B6/J mice was obtained by cardiac puncture and deposited in heparin-coated tubes (BD Microtainer with PST additive). Mice were maintained at Stanford University in compliance with the Stanford University Institutional Animal Care and Use Committee regulations and procedures were approved by the Stanford University Administrate Panel on Laboratory Animal Care. Blood from healthy human donors was obtained by venous blood draw and deposited in heparin-coated tubes (BD Vacutainer with spray-coated sodium heparin additive). The blood was centrifuged at 2000 rcf for 20 minutes and plasma was isolated from the top layer. All work was performed in accordance with the protocol approved by the Stanford IRB Administrative Panel on Human Subjects Research. Plasma reactions included the following: 80% plasma, PBS (or PBS + 50 mM EDTA), and 5 μM cGAMP with trace [^32^P]cGAMP (prepared as described previously^13^). For IC_50_ value assays, 5-fold dilutions of inhibitor in PBS were also included. At indicated times, reaction was quenched by spotting on a HP-TLC silica gel plate (EMD Millipore). The TLC developing solvent was 5 mM ammonium bicarbonate in 85% ethanol/15% water. Plates were exposed on a phosphor screen (GE BAS IP MS), imaged on a Typhoon 9400, and quantified with ImageJ. Half-life values were fit with Graphpad Prism 7.03.

### In vitro enzyme assays

Mouse ENPP1 (3 nM; expressed and purified as described previously^24^) was incubated with 5 μM cGAMP (synthesized as described previously^13^) and 5-fold serial dilutions of compounds in buffer containing 50 mM Tris pH 7.4, 250 nM NaCl, 500 uM CaCl_2_, and 1 uM ZnCl_2_ (total reaction volume = 10 μL). To determine the protein-shifted potency, 40 μM human serum albumin (Sigma) was included in the reaction. Reactions were incubated at room temperature for 3 hours and then heat inactivated at 95°C for 10 minutes. The AMP degradation product was converted to ATP using an enzyme mixture of polyphosphate:AMP phosphotransferase (PAP) and myokinase (MilliporeSigma), which was detected using luciferase (CellTiterGlo, Promega) as previously described (Mardjuki et al., *JBC*, accepted). Briefly, PAP was diluted to 2 mg/mL in buffer containing 50 mM Tris pH 7.5, 0.1% NP-40. Myokinase was diluted to 2 KU/mL in buffer containing 3.2 mM ammonium sulfate pH 6.0, 1 mM EDTA, and 4 mM polyphosphate. The heat-inactivated ENPP1 reaction was incubated with PAP (0.01 μg/μL) and myokinase (0.0075 U/μL) in buffer containing 40 mM Tris pH 7.5, 0.05 mg/mL Prionex, 5 mM Mg Cl_2_, 20 μM polyphosphate, and 0.15 g/L phenol red (for ease of pipetting) for 3 hours (total reaction volume = 20 μL). CellTiterGlo (20 μL) was added to the reaction according to manufacturer’s protocol and luminescence was measured. Data were normalized to 100% enzyme activity (no compound) and 0% enzyme activity (no enzyme) before being fit to the function 100 / (1 + ([compound] / IC_50_)). *K*_i_ values were calculated according to the Cheng-Prusoff equation *K*_i_ = IC_50_ / (1 + [substrate] / *K*_m_). *K*_m_ values were determined by incubating increasing concentrations of cGAMP with trace [^32^P]cGAMP and ENPP1 and measuring initial rates of reaction by TLC and autoradiography (described above in “Plasma kinetics”). Data were plotted as substrate concentration vs. initial velocity (Michaelis-Menten plot) and fit for *K*_m_ (*K*_m_ = 240 μM for mouse ENPP1, substrate cGAMP, at pH 7.4) with Graphpad Prism 7.03.

### Cell viability

Primary human peripheral blood mononuclear cells (PBMCs) were isolated by subjecting enriched buffy coat from whole blood (Stanford Blood Center) to a Percoll density gradient and cultured in RPMI (Cellgro) supplemented with 10% heat-inactivated RBS (Atlanta Biologicals), 100 U/mL penicillin-streptomycin (ThermoFisher), and 50 μM 2-mercaptoethanol (Sigma) and 10 ng/mL M-CSF (PeproTech). Cells were maintained in a 5% CO_2_ incubator at 37°C. Cells were plated in 384 well plates at 2000 cells/well incubated with 5-fold dilutions of compounds for 16 hours. Viability was measured with CellTiterGlo (Promega) and normalized to 100% viability (no compound) in Graphpad Prism 7.03.

### Determination of the structure of mouse ENPP1 with compound 15

The extracellular region (residues 92–905) of mouse ENPP1 was expressed, purified, and crystallized as described previously.^24^ Crystals of ENPP1 in complex with compound 15 were grown at 20 °C by the hanging-drop vapor-diffusion method by mixing 1 μL of the protein solution [5 mg mL−1 in 5 mM Tris·HCl (pH 8.0), 150 mM NaCl, and 0.2 mM ZnSO_4_] and 1 μL of the reservoir solution [17–19% (v/v) PEG600, 50 mM sodium acetate (pH 4.5), 50 mM magnesium acetate, 0.5% (w/v) polyvinylpyrrolidone and 1 mM compound 15]. Crystals were harvested and cryo-cooled in liquid nitrogen.

Multiple needle-like crystals were tested for X-ray diffraction which, when X-ray exposed, showed weak diffracting power. One needle crystal was isolated and used to collect a data set to a minimum d-spacing of around 3.2 Å. Data was collected at cryogenic temperature (100 K) at beamline 5.0.1 of the Advanced Light Source (ALS) Synchrotron (Berkeley, CA, USA) at a single 0.97741 Å wavelength. Data was reduced with Mosflm^44^ and scaled with SCALA^45^ within the CCP4 suite.^46^ The crystal belonged to the trigonal space group P 3_1_ and contained two polypeptide chains per asymmetry unit. Data collection and refinement statistics are listed in Table 3. The structure was solved by the molecular replacement method with Phaser^47^ using mouse ENPP1 (PDB code: 4GTW) polypeptide chain stripped from ligands, ions, and glycan chains, as the search model. Structural refinement was done using REFMAC^48^ iteratively with visual inspection of electron density maps and manual adjustment of atomic coordinates in COOT^49^ until progression to convergence. The final refined structure shows an excellent agreement with reference protein data as shown by Ramachandran statistics (Table 4). Data collection statistics are derived from SCALA.^45^ To calculate *R*_free_, 5% of the reflections were excluded from the refinement. *R*_sym_ is defined as *R*_sym_ = Σ_hkl_Σ_i_|I_i_(hkl) - <I(hkl)>| / Σ_hk_lΣ_i_I_i_(hkl). Data refinement statistics are derived from REFMAC.^48^ The final quality check was done with Procheck.^50^ Graphic renderings were prepared with Pymol.^51^

As previously observed,^24^ the structure could be traced unambiguously from residue 170 to 902. No electron density is observed for the two SMB-like domains (residues 92-169). The two independent polypeptide chains in the asymmetric unit consist of a catalytic domain (residues 190–578), a nuclease-like domain (residues 629–902), and two long linker regions, L1 and L2 (residues 170–189 and 579–628, respectively). Portions of the structure not seen in the electron density map and not modeled belong predominantly to the nuclease-like domain and linker L2 region: chain A, residues 506-512; 609-628; 681-689; 708-715; 725-735; 799-803; 848-855, and chain B, residues 507-511; 607-629; 681-692; 709-715; 726-734; and 848-855. The structure has N-glycosylated sites at Asn267, Asn323, and Asn567. The two zinc ions are bound within the active site of ENPP1 and coordinated by Asp358, His362, and His517 at one site, and by Asp200, Thr238, Asp405, and His406 at the other. In the nuclease domain, a calcium ion is coordinated by the side chains of Asp780, Asp782, Asp784, and Asp788 and the main chain carbonyl group of Arg786, forming an EF hand-like motif. Extra electron density unaccounted for by the model allowed placement of one copy of compound 15 in the substrate-binding pocket with the dimethoxyquinazoline ring sandwiched between the side chains of Phe239 and Tyr322 and the phosphonic acid group between the zinc atoms. In the final rounds of refinement, the ligand was modeled at 0.5 occupancy in the binding pocket of chain A. The very weak electron density of the cycloalkyl carbon atoms prevented to accurately model its pucker and therefore the piperidine ring was tentatively assigned a chair shape with the nitrogen and the 4-carbon lying out-of-plane. This seemed to be a reasonable choice as a survey of the Cambridge Structural Database^52^ revealed that in three available quinazolinylpiperidinyl heterocycle crystal structures (CCDC codes: LUMVIA, PASNOP, ZAYQIC) the piperidine ring is in the chair conformation.

### Mouse pharmacokinetics

Mice were injected via SC or retro-orbital IV route with 10 mg/kg of compound 32 (200 μL of 2 mg/mL dissolved in PBS). At indicated time points, mice were sacrificed and blood, kidneys, and livers were collected. Alternatively, mice were injected via SC with 300 mg/kg of compound 32 (200 μL of 60 mg/mL dissolved in PBS). At indicated time points, 50-100 μL of blood was collected from live mice via cheek puncture. Blood was allowed to clot at room temperature for 30 minutes before centrifuging at 1000xg to isolate serum. Organs were homogenized with 2.0 mm disruption beads (RPI) in a tissue homogenizer (Bertin Precellys 24) in PBS. Organ homogenates and serum were precipitated with acetonitrile, centrifuged at 16,000xg, and resuspended in a matrix of 2:1 0.1% formic acid:acetonitrile with clemizole as the internal standard. LC-MS/MS analysis was performed on a Shimadzu HPLC with an autosampler set at 4 °C and connected to an AB Sciex 4000 QTRAP.

### Mouse tumor models

Five- to nine-week-old female C57BL6/J mice (Jackson Laboratories) were inoculated with 5 × 10^4^ E0771 cells suspended in 50 μL of PBS in the fifth mammary fat pad. When tumor volume (determined by length^2^ × width/2) reached 100 ± 20 mm^3^, mice were injected via SC route with 300 mg/kg of compound 32 as described above, once daily for seven consecutive days. Tumor volumes were recorded up to the humane endpoint (tumor volume exceeding 1000 mm^3^), when mice were sacrificed. Animal death was plotted with GraphPad Prism 7.03, and statistical significance was assessed by the Gehan-Breslow-Wilcoxon test. Mice were maintained at Stanford University in compliance with Stanford University Institutional Animal Care and Use Committee regulations and procedures were approved by the Stanford University administrate panel on laboratory animal care.

## Figures

**Figure S1.**
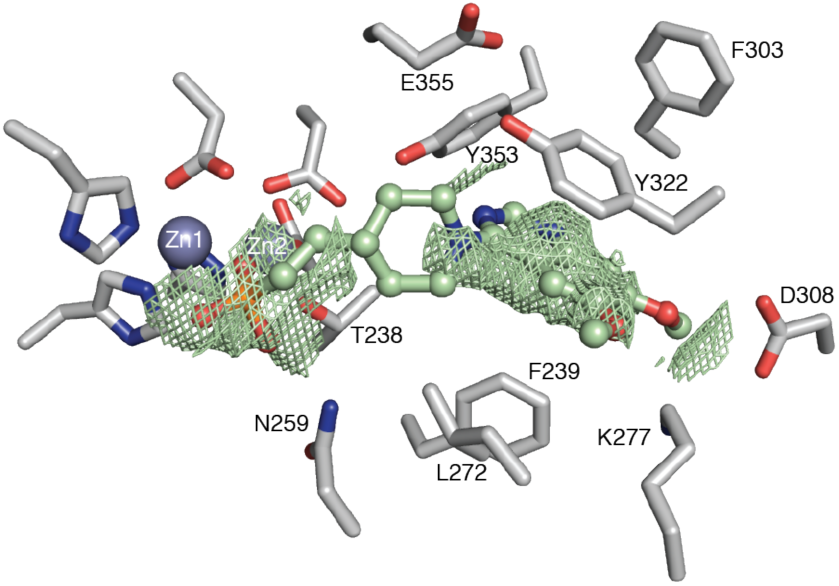
Co-crystal structure of ENPP1 and compound 15 with electron density map. σ-weighted 2Fo-Fc map (0.5 σ; green mesh) around compound **15** (green sticks) in complex with ENPP1 (gray sticks).

**Figure S2.**
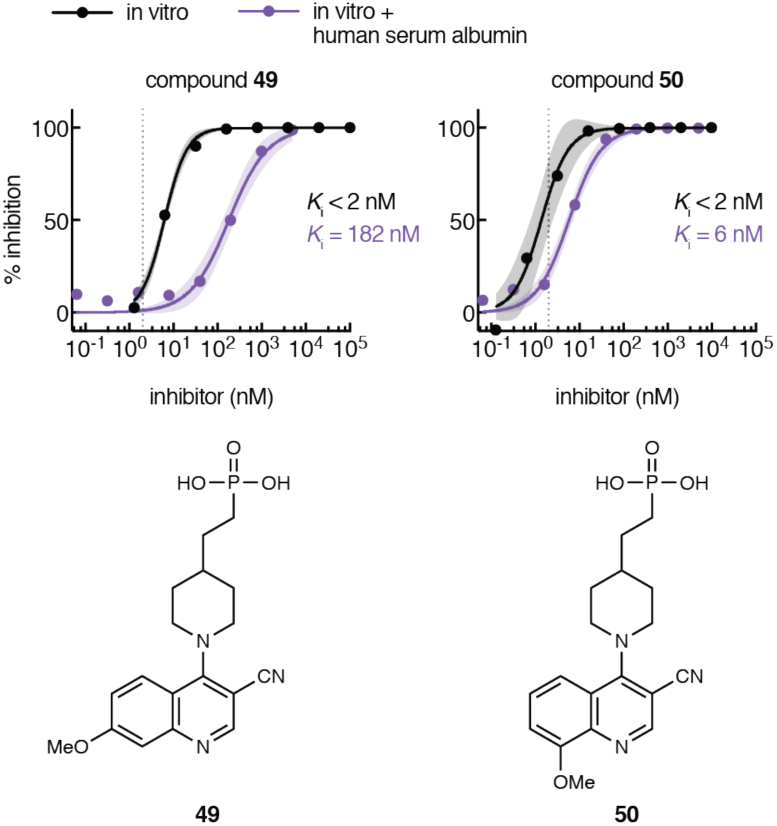
Protein-shifted *K*_i_ values for selected piperidine-3-nitrile-quinoline compounds. In vitro dose-inhibition curves for selected top ENPP1 inhibitors. *K*_i_ values were determined using 3 nM ENPP1 and 5 μM cGAMP and (where indicated) in the presence of 40 μM human serum albumin at pH 7.4. Dotted line represents the minimum IC_50_ value (2 nM) measurable with the given enzyme concentration. Chemical structures of inhibitors are displayed below. Dots represent the mean of two independent replicates, and shaded areas around the fitted curves represent the 95% confidence interval of the fit.

**Figure S3.**
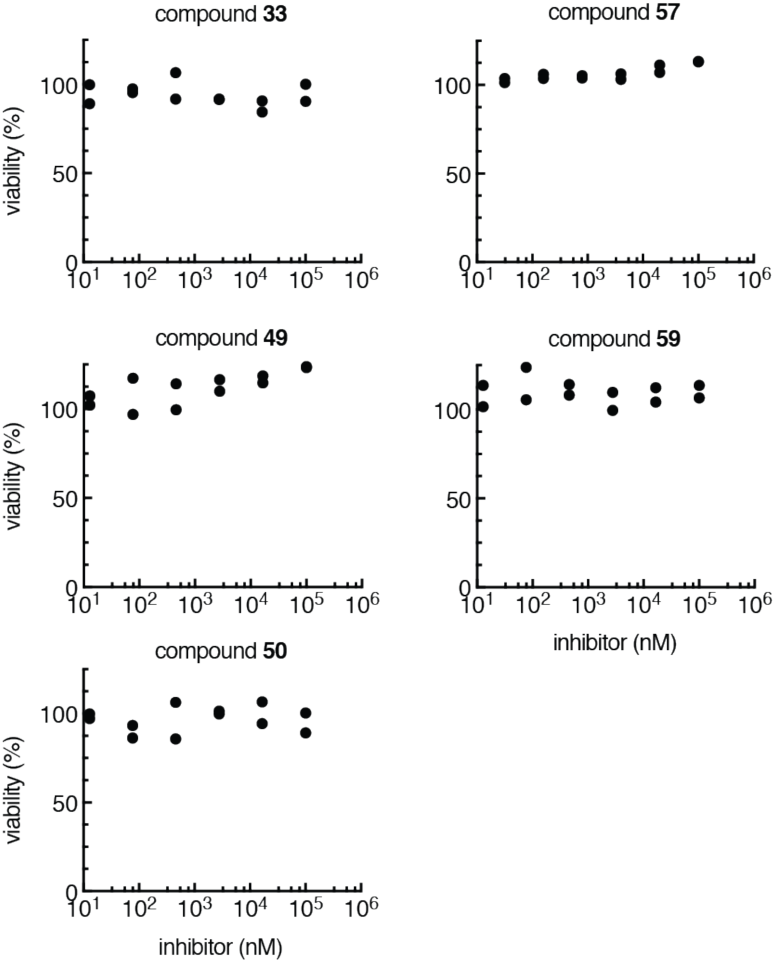
Cell viability after treatment with selected ENPP1 inhibitors. Cell viability of primary human peripheral blood mononuclear cells (PBMCs) after incubation with indicated compounds for 16 hours measured by CellTiterGlo. Data is normalized to no compound (100% cell viability). Two cell culture replicates are plotted.

## Supplemental Tables

**Table S1:** Intermolecular interactions in the co-crystal structure of compound 15 with ENPP1.

| Interacting partners |  | Distance (Å) | Type |
| --- | --- | --- | --- |
| ENPP1 | Compound 15 |  |  |
| Asp308 OD1, OD2 | O14 (7-methoxy) | 3.43, 2.93 | H-bond |
| Lys277 NZ | O14 (7-methoxy) | 3.42 | H-bond |
| Asn259 ND | O25 (phosphonate) | 2.96 | H-Bond |
| Zinc | O26, O27<br>(phosphonate) | 2.08, 2.14, 2.51 | Metal coordination |
| L272 | Piperidine ring | 3.5–3.8 | Hydrophobic |
| Tyr322 | Quinazoline ring | 3.77 (average), range:<br>3.31–4.50 | Aromatic (sandwich<br>stacking) |
| Phe239 | Quinazoline ring | 3.76 (average) | Aromatic (T-shaped<br>stacking) |
| Tyr353 | Quinazoline ring | 2.88–3.44 | Hydrophobic |
| Tyr353 OH | Quinazoline ring N3 | 3.1 | Polar |

**Table S2:** Data collection and refinement statistics.

| ENPP1-compound 15 |  |
| --- | --- |
| **Data collection** |  |
| Space group | P3<sub>1</sub> |
| Cell dimensions |  |
| *a*, *b*, *c* (Å) | 102.35, 102.35, 172.89 |
| $\alpha$, $\beta$, $\gamma$ (°) | 90, 90, 120 |
| Resolution (Å) | 88.64–3.20 (3.37–3.20) |
| *R*<sub>sym</sub> or *R*<sub>merge</sub> | 0.160 (0.723) |
| *I* / σ*I* | 5.4 (1.8) |
| Completeness (%) | 91.8 (90.6) |
| Redundancy | 2.1 (2.1) |
| **Refinement** |  |
| Resolution (Å) | 30.64–3.20 |
| No. reflections | 29,094 |
| *R*<sub>work</sub> / *R*<sub>free</sub> | 20.1/27.5 |
| No. atoms | 10,874 |
| Protein | 10,682 |
| Ligand/ion/other | 26 (1x compound 15), 6 ions (4x<br>Zn, 2x Ca), 128 glycan (6x NAG,<br>4x mannose) |
| Water | 32 |
| *B*-factors |  |
| Protein | 66.5 |
| Ligand/ion/other | 81.8 (compound 15), 60.1 (ions),<br>86.8 (glycan) |
| Water | 35.5 |
| R.m.s. deviations |  |
| Bond lengths (Å) | 0.009 |
| Bond angles (°) | 1.441 |

Values in parentheses are for highest-resolution shell. Data collection statistics are derived from SCALA.<sup>45</sup> To calculate *R*<sub>free</sub>, 5% of the reflections were excluded from the refinement. *R*<sub>sym</sub> is defined as $R_{\text{sym}} = \sum_{\text{hkl}} \sum_i |I_i(\text{hkl}) - \langle I(\text{hkl}) \rangle| / \sum_{\text{hkl}} \sum_i I_i(\text{hkl})$ . Data refinement statistics are derived from REFMAC. The final quality check was done with Procheck.

**Table S3.**
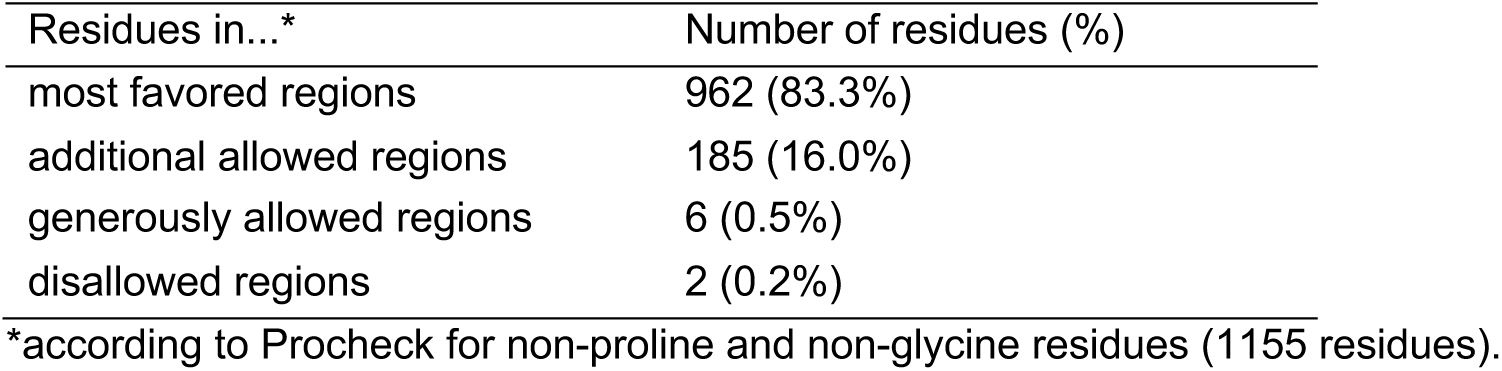
Ramachandran statistics.

