## Supplemental Methods for "Structure-aided development of small molecule inhibitors of ENPP1, the extracellular phosphodiesterase of the immunotransmitter cGAMP"

**Chemical Synthesis:** Reactions were performed under ambient atmosphere unless otherwise noted. Qualitative TLC analysis was performed on 250 mm thick, 60 Å, glass backed, F254 silica (Silicycle, Quebec City, Canada). Visualization was accomplished with UV light and exposure to *p*-anisaldehyde or $KMnO_4$ stain solutions followed by heating. All solvents used were ACS grade Sure-Seal, and all other reagents were used as received unless otherwise noted. Flash chromatography was performed on a Teledyne Isco purification system using silica gel flash cartridges (SiliCycle®, SiliaSep™ 40-63 µm, 60Å). HPLC was performed on an Agilent 1260 Infinity preparative scale purification system using an Agilent PrepHT Zorbax Eclipse XDB-C18 reverse-phase column (21.2 × 250 mm). Structure determination was performed using $^1H$ spectra that were recorded on a Bruker AV-500 spectrometer, and low-resolution mass spectra (ESI-MS) that were collected on a Shimadzu 20-20 ESI LCMS instrument. Structure determination was performed using $^1H$ spectra that were recorded on either a Bruker AV-500 or AV-400 spectrometer, and low-resolution mass spectra (ESI-MS) that were collected on a Shimadzu 20-20 ESI LCMS instrument. Final compound purity was >95%, as determined by HPLC-MS. All final compound $^1H$ spectra were consistent with the expected structures.

**Preparation of 2-((6-amino-9*H*-purin-8-yl)thio)-*N*-(3,4-dimethoxyphenyl)acetamide 1.**

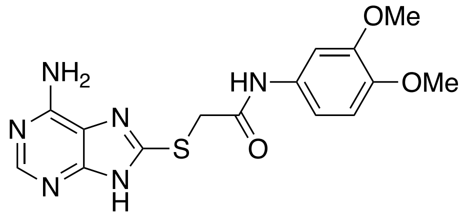


Compound **1** was prepared according to Chang, et al. *J. Med. Chem.* 2014, 57, 10080−10100.

LCMS: [M − H]$^-$ $m/z$ 359.1 $^1$H NMR (500 MHz, DMSO-*d6*) δ 10.73 (br s, 1H), 8.02 (s, 1H), 7.25 (s, 1H), 7.10 (d, $J$ = 8.6 Hz, 1H), 6.88–6.84 (m, 3H), 4.06 (s, 2H), 3.70 (s, 6H),

**Synthesis of the ureas 4 and 5.**

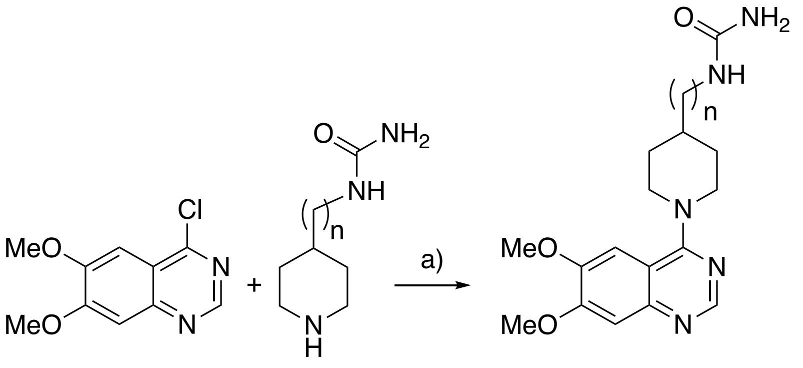


a) DIPEA, i-PrOH.

**Preparation of 1-(2-(1-(6,7-dimethoxyquinazolin-4-yl)piperidin-4-yl)ethyl)urea 4**

To a solution of 1-(2-(piperidin-4-yl)ethyl)urea **64** (173 mg, 1.01 mmol) in isopropanol (5 mL) was added 4-chloro-6,7-dimethoxyquinazoline **63** (181 mg, 0.81 mmol) and *N,N*-diisopropylethylamine (391 mg, 3.03 mmol) under nitrogen atmosphere. The mixture was stirred at room temperature for 2 h and then evaporated to dryness under reduced pressure. Purification (prep-HPLC) gave the title compound **4** (172 mg, 47%) as light yellow crystals.

LCMS: [M + H]$^+$ $m/z$ 360. $^1$H NMR (400 MHz, DMSO-$d_6$) δ 8.49 (s, 1H), 7.17 (s, 1H), 7.07 (s, 1H), 5.92–5.90 (m, 1H), 5.36 (br s, 2H), 4.13–4.09 (m, 2H), 3.90 (s, 3H), 3.88 (s, 3H), 3.04–2.94 (m, 4H), 1.81–1.78 (m, 2H), 1.62–1.56 (m, 1H) and 1.38–1.33 (m, 4H).

**Preparation of 1-((1-(6,7-dimethoxyquinazolin-4-yl)piperidin-4-yl)methyl)urea 5.**

To a solution of 1-(piperidin-4-ylmethyl)urea **65** (155 mg, 0.97 mmol) in isopropanol (10 mL) was added 4-chloro-6,7-dimethoxyquinazoline **63** (174 mg, 0.78 mmol) and *N,N*-diisopropylethylamine (394 mg, 2.9 mmol) under a nitrogen atmosphere. The mixture was stirred at 10 ºC for 3 h and then evaporated to dryness under reduced pressure. Chromatography ($SiO_2$: 0 to 6% MeOH in dichloromethane) to give the desired product **5** (150 mg, 44%) as a white solid. LCMS: $[M + H]^+$ *m/z* 346.0 $^1H$ NMR (400 MHz, Methanol-$d_4$) δ 8.44 (s, 1H), 7.14 (s, 1H), 7.12 (s, 1H), 4.28–4.24 (m, 2H), 3.96 (s, 3H), 3.94 (s, 3H), 3.13–3.07 (m, 4H), 1.94–1.87 (m, 3H) and 1.50–1.41 (m, 2H).

**Preparation of 3-(1-(6,7-dimethoxyquinazolin-4-yl)piperidin-4-yl)propanoic acid 6.**

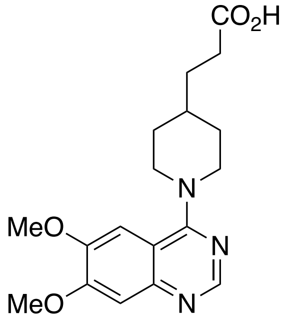


4-Chloro-6,7-dimethoxy-quinazoline **63** (3.14g, 13.98 mmol) and 3-(4-piperidyl)propanoic acid (2.0 g, 12.72 mmol) were suspended in isopropanol (100 mL) and stirred at 90 °C for 3 h. Once cooled, the mixture was evaporated to dryness under reduced pressure. The residue was then triturated with $CH_2Cl_2$ (20 mL) to give the title compound **6** (1.87 g, 42%) as a white solid. LCMS: $[M + H]^+$ *m/z* 346.1 $^1H$ NMR (500 MHz, DMSO-$d_6$) δ 12.12 (br s), 8.54 (s, 1H), 7.23 (s, 1H), 7.14 (s, 1H), 4.18 (d, *J* = 10.4 Hz, 2H), 3.96 (s, 3H), 3.94 (s, 3H), 3.04 (t, *J* = 12.5 Hz, 2H), 2.27 (t, *J* = 7.5 Hz, 3H), 1.44–1.39 (m, 6H).

**Preparation of (2-(1-(6,7-dimethoxyquinolin-4-yl)piperidin-4-yl)ethyl)boronic acid 7.**

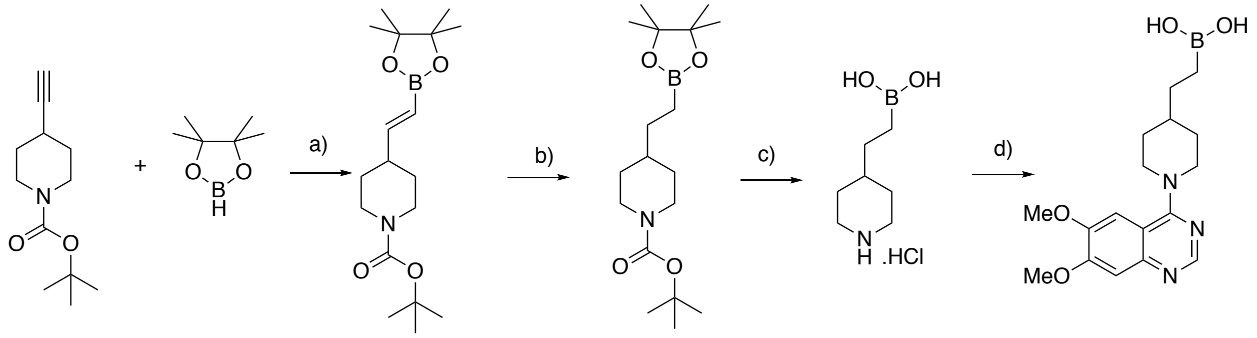

66 67 68 69 70 7

a) $Cp_2ZrCl_2$, PhMe 60 °C; b) Pd/C, $H_2$, MeOH; c) HCl (aq), MeOH/Hexanes; d) **63**, DIEA, THF 80 °C.

A solution of *tert*-butyl 4-ethynylpiperidine-1-carboxylate **66** (2.92 g, 13.95 mmol), bis(cyclopentadienyl)zirconium chloride hydride (150 mg, 0.518 mmol) and 4,4,5,5-tetramethyl-1,3,2-dioxaborolane **67** (1.49 g, 11.63 mmol) in toluene (100 mL) was stirred at 60 ºC for 16 h and then diluted with ether and evaporated to dryness under reduced pressure. Chromatography ($SiO_2$; 2–5% ethyl acetate in petroleum ether) provided **68** (4.2 g, 89 %). To a mixture of tert-butyl (*E*)-4-(2-(4,4,5,5-tetramethyl-1,3,2-dioxaborolan-2-yl)vinyl)piperidine-1-carboxylate **68** (4.2 g, 12.46 mmol) and palladium on carbon (840 mg, 20% w/w) in MeOH (500 mL) was placed under an atmosphere of hydrogen and stirred at room temperature for 16 h. The mixture was then filtered through a pad of Celite® and then evaporated to dryness under reduced pressure to afford **69** (4.2 g, 92 %). 1M Aqueous HCl solution (4 mL) solution was added to a cooled (0 ºC) mixture of **73** (460 mg, 1.36 mmol) in MeOH/hexane (5 mL/5 mL). The mixture was allowed to warm to room temperature and stirred for 3 h and then evaporated to dryness under reduced pressure to afford (2-(piperidin-4-yl)ethyl)boronic acid **70** (180 mg, 68 %) as the hydrochloride salt. To a solution of **70** (140 mg, 1.04 mmol) in THF (5 mL) was added 4-chloro-6,7-dimethoxyquinazoline **63** (180 mg, 0.935 mmol) followed by *N,N*-diisopropylethylamine (360 mg, 1.87 mmol). The mixture was stirred at 80 °C for 16 h and then evaporated to dryness under reduced pressure. Purification (prep-HPLC) gave the title compound as a light yellow solid (105 mg; 37%).

LCMS: $[M + H]^+$ *m/z* 346.3. $^1H$ NMR (400 MHz, DMSO-$d_6$) δ 8.67 (s, 1H), 7.26 (s, 1H), 7.25 (s, 1H), 4.62–4.59 (m, 2H), 3.92 (s, 3H), 3.90 (s, 3H), 3.42–3.36 (m, 4H), 2.46 (s, 1H), 1.88–1.86 (m, 2h), 1.29–1.14 (m, 3H) and 0.60–0.56 (m, 2H).

**Preparation of the hydroxamic acids 8 and 9.**

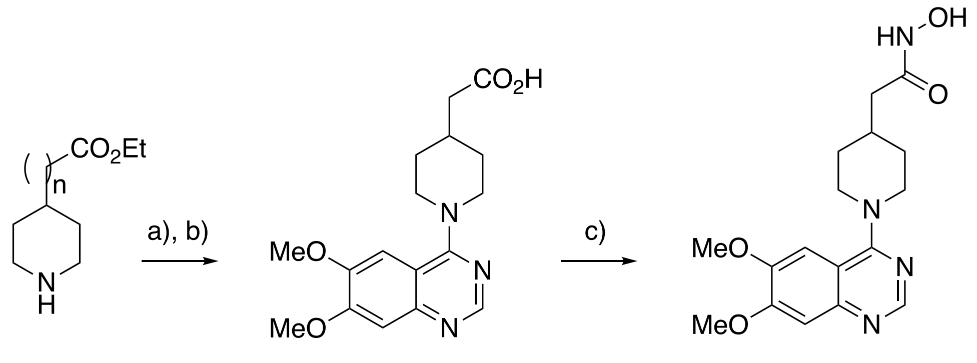


**71** n = 1
**73** n = 0

**72** n = 1
**74** n = 0

**8** n = 1
**9** n = 0

a) **63**, i-PrOH, 100 °C; b) NaOH, THF, $H_2O$; c) $NH_2OH.HCl$, DIPEA, BOP, THF.

**Preparation of 2-(1-(6,7-dimethoxyquinazolin-4-yl)piperidin-4-yl)-*N*-hydroxyacetamide 8.**

A mixture of 4-chloro-6,7-dimethoxyquinazoline **63** (600 mg, 2.68 mmol) and ethyl 2-(piperidin-4-yl)acetate **71** (504 mg, 2.95 mmol) in *i*-PrOH (6 mL) was stirred at 100 °C for 16 h in a sealed tube. Then the reaction mixture was concentrated under reduced pressure and the residue was purified by silica gel chromatography to give ethyl 2-(1-(6,7-dimethoxyquinazolin-4-yl)piperidin-4-yl)acetate (750 mg, 77%). 2M NaOH solution in $H_2O$ (1 mL) was added to a mixture of ethyl 2-(1-(6,7-dimethoxyquinazolin-4-yl)piperidin-4-yl)acetate (250 mg, 0.696 mmol) in THF (10 mL). The mixture was stirred at room temperature for 16 h and then quenched by the addition of 1 M HCl solution. The organic phase was extracted with ethyl acetate, washed with brine, dried ($Na_2SO_4$) and evaporated to dryness under reduced pressure to give the acid **72** (200 mg, 86%) as a white solid.

To a mixture of the acid **72** (300 mg, 0.906 mmol) in THF (10 mL) was added $NH_2OH{\cdot}HCl$ (76 mg, 1.09 mmol), DIEA (468 mg, 3.63 mmol) and (benzotriazole-1-yloxy)tris(dimethylamino)phosphonium hexafluorophosphate (BOP) (481 mg, 1.09 mmol). The mixture was stirred at room temperature for 16 h and then diluted with water, extracted with ethyl acetate, washed with brine solution, dried ($Na_2SO_4$) and evaporated to dryness under reduced pressure. Chromatography ($SiO_2$, 0-100% ethyl acetate in petroleum ether) to give the title product **8** (180 mg, 77%) as a white solid. LCMS: $[M + H]^+$ *m/z* 347.10.$^1$H NMR (400 MHz, $D_2O$) δ 8.42 (s, 1H), 7.13 (s, 1H), 7.00 (s, 1H), 4.68–4.62 (m, 2H), 3.95 (s, 3H), 3.91 (s, 3H), 3.51–3.45 (m, 2H), 2.21–2.15 (m, 3H), 1.93–1.0 (m, 2H) and 1.45–1.36 (m, 2H).

**Preparation of 1-(6,7-dimethoxyquinazolin-4-yl)-*N*-hydroxypiperidine-4-carboxamide 9.**

Was synthesized according to the procedure for **8** but using ethyl piperidine-4-carboxylate **73**.

LCMS: $[M + H]^+$ *m/z* 333.25.$^1$H NMR (400 MHz, $D_2O$) δ 8.39 (s, 1H), 7.04 (s, 1H), 6.94 (s, 1H), 4.62–4.58 (m, 2H), 3.91 (s, 3H), 3.86 (s, 3H), 3.47–3.41 (m, 2H), 2.65–2.60 (m, 1H), 1.97–1.94 (m, 2H) and 1.82–1.77 (m, 2H).

### General Procedure for compounds 10, 11, 12, 13 and 16.

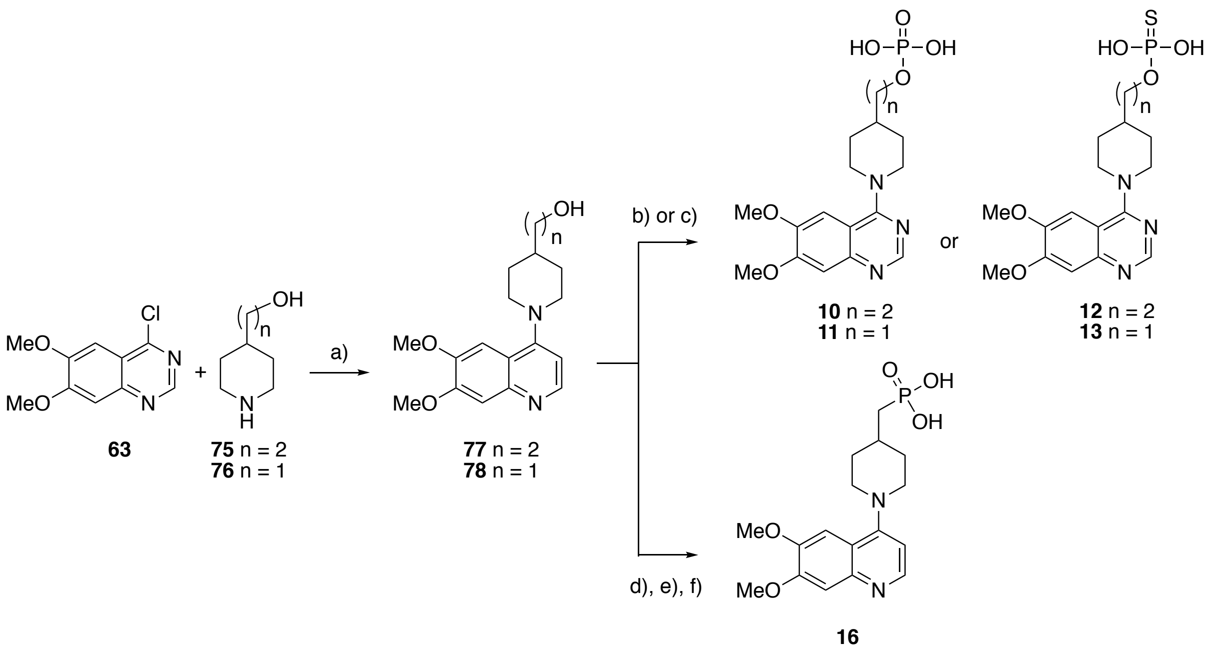

a) i-PrOH, 100 °C; b) pyridine, $PSCl_3$ then $H_2O$; or c) pyridine, $POCl_3$, then $H_2O$; d) $PPh_3$, $I_2$, imidazole, $CH_2Cl_2$; e) $PO(OBn)_2$, DBU, MeCN; f) Pd/C, $H_2$, MeOH.

### Preparation of 2-(1-(6,7-dimethoxyquinolin-4-yl)piperidin-4-yl)ethan-1-ol 77.

A mixture of 4-chloro-6,7-dimethoxyquinazoline **63** (1.0 g, 4.46 mmol) and piperidin-4-ylethanol **79** (633 mg, 4.91 mmol) in isopropanol (10 mL) was stirred at 100 °C for 16 h in a sealed tube. Upon cooling, the reaction mixture was concentrated under reduced pressure and the residue was purified by silica gel chromatography ($SiO_2$; EtOAc in petroleum ether) to give 2-(1-(6,7-dimethoxyquinolin-4-yl)piperidin-4-yl)ethan-1-ol **75** (1.3 g, 91%).

### Preparation of (1-(6,7-dimethoxyquinolin-4-yl)piperidin-4-yl)methanol 78.

A mixture of 4-chloro-6,7-dimethoxyquinazoline **63** (900 mg, 4.02 mmol) and piperdin-4-ylmethanol **76** (508 mg, 4.42 mmol) in *i*-PrOH (10 mL) was stirred at 100 °C for 16 h in a sealed tube. Upon cooling, the reaction mixture was evaporated to dryness under reduced pressure.

Purification by chromatography ($SiO_2$; 10 to 80% ethyl acetate in petroleum ether) to give (1-(6,7-dimethoxyquinolin-4-yl)piperidin-4-yl)methanol **78** (1 g, 82%).

**Preparation of 2-(1-(6,7-dimethoxyquinazolin-4-yl)piperidin-4-yl)ethyl dihydrogen phosphate 10.**

2-(1-(6,7-Dimethoxyquinazolin-4-yl)piperidin-4-yl)ethan-1-ol **77** (340 mg, 1.07 mmol) was dissolved in 10 mL dry pyridine, then it was cooled to −15 °C and stirred for 10 min. $POCl_3$ (821 mg, 5.4 mmol) was added dropwise under a $N_2$ atmosphere, The reaction temperature was raised to 0 °C slowly, then stirred for another 30 min again. The mixture was poured into sodium hydrogen carbonate solution (800 mg in 250 mL water) at 0 °C. The desired compound was extracted with dichloromethane. The organic phase was concentrated and purified with Prep-HPLC to give 2-(1-(6,7-dimethoxyquinazolin-4-yl)piperidin-4-yl)ethyl dihydrogen phosphate **10** (52 mg, 12%) as a white solid.

LCMS: $[M + H]^+$ *m/z* 398. $^1H$ NMR (400 MHz, DMSO-$d_6$) δ 8.54 (s, 1H), 7.17 (s, 1H), 7.15 (s, 1H), 4.28–4.16 (m, 2H), 3.93 (s, 8H), 3.13–3.04 (m, 2H), 1.90–1.80 (m, 2H), 1.75 (s, 1H), 1.59 (d, *J* = 6.4 Hz, 2H) and 1.44–1.32 (m, 2H).

**Preparation of (1-(6,7-dimethoxyquinolin-4-yl)piperidin-4-yl)methyl dihydrogen phosphate 11.**

(1-(6,7-Dimethoxyquinolin-4-yl)piperidin-4-yl)methanol **78** (100 mg, 0.33 mmol) was dissolved in dry pyridine (3 mL), then it was cooled to −15 ℃ and stirred for 10 min. $POCl_3$ (253 mg, 1.65 mmol) was added dropwise under nitrogen atmosphere. The reaction temperature was raised to 0 °C slowly, then stirred for another 30 min. The mixture was poured into aqueous $NaHCO_3$ solution (160 mg in 50 mL of water) at 0 °C. The desired compound was extracted with dichloromethane and then evaporated to dryness under reduced pressure. Purification by Prep-HPLC afforded (1-(6,7-dimethoxyquinolin-4-yl)piperidin-4-yl)methyl dihydrogen phosphate **11** (70 mg, 55%) as white powder after lyophilization. LCMS: $[M + H]^+$ *m/z* 384.20. $^1H$ NMR (400 MHz, DMSO-$d_6$) δ 8.74 (d, *J* = 1.7 Hz, 1H), 7.31 (s, 1H), 7.20 (s, 1H), 4.66 (d, *J* = 13.0 Hz, 1H), 3.97 (m, J = 12.6, 1.6 Hz, 8H), 3.76 (t, *J* = 6.6 Hz, 3H), 2.19–2.00 (m, 1H), 1.92 (d, *J* = 13.5 Hz, 2H), 1.45 (dd, *J* = 14.2, 10.7 Hz, 1H).

**Preparation of *O*-(2-(1-(6,7-dimethoxyquinazolin-4-yl)piperidin-4-yl)ethyl) *O,O*-dihydrogen phosphorothioate 12.**

To a solution of 2-(1-(6,7-dimethoxyquinolin-4-yl)piperidin-4-yl)ethan-1-ol **77** (150 mg, 0.473 mmol) in dry pyridine (5 mL) was added $P(S)Cl_3$ (477 mg, 2.84 mmol) at –15 °C. After being stirred at 0 °C for 0.5 h, the mixture was poured over a solution of $NaHCO_3$ (238 mg, 2.84 mmol) in $H_2O$ (50 mL). The mixture was stirred at 0 °C for 2 h. The progress of the reaction mixture was monitored by LCMS. Then the mixture was concentrated under reduced pressure and the residue was purified by prep-HPLC to afford compound **12** (16 mg, 8%) as a light yellow solid. LCMS: $[M + H]^+$ *m/z* 414.05. $^1H$ NMR (400 MHz, DMSO-$d_6$) δ 8.62 (s, 1H), 7.19 (d, *J* = 7.7 Hz, 2H), 4.45 (d, *J* = 12.3 Hz, 2H), 3.91 (d, *J* = 11.3 Hz, 10H), 1.86 (d, *J* = 12.2 Hz, 3H), 1.56 (d, *J* = 6.4 Hz, 2H), 1.34 (d, *J* = 10.7 Hz, 2H).

**Preparation of *O*-((1-(6,7-dimethoxyquinazolin-4-yl)piperidin-4-yl)methyl)*O,O*-dihydrogen phosphorothioate 13.**

To a solution of (1-(6,7-dimethoxyquinolin-4-yl)piperidin-4-yl)methanol **78** (100 mg, 0.330 mmol) in dry pyridine (5 mL) was added $P(S)Cl_3$ (280 mg, 1.98 mmol) at –15 °C. After being stirred at 0 °C for 0.5 h, the mixture was poured over a solution of $NaHCO_3$ (116 mg, 1.98 mmol) in $H_2O$ (50 mL). The mixture was stirred at 0 °C for 2 h. The mixture was evaporated to dryness under reduced pressure and the residue was purified by prep-HPLC to afford compound **13** (10 mg, 7.6%) as a yellow solid. LCMS: $[M + H]^+$ *m/z* 400.15. $^1H$ NMR (400 MHz, DMSO-$d_6$) δ 8.54 (s, 1H), 7.18 (s, 1H), 7.11 (s, 1H), 4.25 (d, *J* = 13.4 Hz, 2H), 3.89 (d, *J* = 9.1 Hz, 6H), 3.76 (s, 2H), 3.10 (d, *J* = 11.8 Hz, 3H), 1.94 (s, 1H), 1.81 (d, *J* = 12.7 Hz, 2H), 1.39 (d, *J* = 11.4 Hz, 1H).

**Preparation of ((1-(6,7-dimethoxyquinazolin-4-yl)piperidin-4-yl)methyl)phosphonic acid 16**.

$PPh_3$ (3.39 g, 15 mmol) and imidazole (1.02 g, 15 mmol) in anhydrous $CH_2Cl_2$ (40 mL) was stirred in 0 °C for 10 min and then $I_2$ (3.8g, 15 mmol) was added. The crude reaction mixture was placed under a nitrogen atmosphere and stirred for a further 10 min and then (1-(6,7-dimethoxyquinazolin-4-yl)piperidin-4-yl)methanol **78** (3.03 g, 10 mmol) was added. The reaction mixture was left to stir at room temperature overnight. The reaction was quenched by the addition of aqueous $Na_2S_2O_3$. The crude mixture was extracted with $CH_2Cl_2$, washed with water, brine, dried ($Na_2SO_4$) and evaporated to dryness under reduced pressure. Recrystallisation from methanol gave 4-(4-(iodomethyl)piperidin-1-yl)-6,7-dimethoxyquinazoline (2.28 g, 56%) as light yellow solid. LCMS: $[M + H]^+$ *m/z* 414.3. $^1H$ NMR (400 MHz, $CDCl_3$) δ 8.63 (d, *J* = 1.3 Hz, 1H), 7.28 (s, 1H), 7.07 (s, 1H), 4.23 (s, 2H), 4.00 (s, 6H), 3.19 (d, *J* = 6.5 Hz, 2H), 3.08 (s, 2H), 2.11–2.00 (m, 2H), 1.82 (s, 1H), 1.49 (s, 2H), 1.29–1.20 (m, 1H). 1,8-Diazabicyclo(5.4.0)undec-7-ene (DBU) (9.2 g, 60.5 mmol) was added to a cooled (0 ºC) solution of bis(benzyloxy)(oxo)-λ4-phosphane (9.5 g, 36.3 mmol) in anhydrous MeCN (40 mL). After 10 min, 4-(4-

(iodomethyl)piperidin-1-yl)-6,7-dimethoxyquinazoline (5.0 g, 12.1 mmol) was added. The resulting mixture was stirred for overnight and then evaporated to dryness under reduced pressure. The residue was dissolved in ethyl acetate, washed with water, brine, dried ($MgSO_4$) and evaporated to dryness under reduced pressure. Purification with FCC [$CH_2Cl_2$:MeOH (50:1)] provided dibenzyl ((1-(6,7-dimethoxyquinazolin-4-yl)piperidin-4-yl)methyl)phosphonate (1.1g, 18%) as a colorless viscous oil. LCMS: $[M + H]^+$ *m/z* 548.20. $^1$H NMR (400 MHz, $CDCl_3$) δ 8.57 (s, 1H), 7.89 (s, 1H), 7.39–7.33 (m, 10H), 6.99 (s, 1H), 5.08 (m, 3H), 4.96 (m, 2H), 4.64 (d, *J* = 13.5 Hz, 2H), 4.09 (s, 3H), 3.93 (s, 3H), 3.27 (d, *J* = 12.9 Hz, 2H), 2.05 (d, *J* = 13.9 Hz, 5H), 1.76 (m, 4H), 1.42 (d, *J* = 12.5 Hz, 2H).

A mixture containing dibenzyl ((1-(6,7-dimethoxyquinazolin-4-yl)piperidin-4-yl)methyl)phosphonate (660 mg, 1.2 mmol) and Pd/C (132 mg, 20% w/w) in MeOH (20 mL) was placed under an atmosphere of $H_2$ and stirred at room temperature. After 4 h, the crude mixture was filtered through a pad of Celite® and the filtrate was evaporated to dryness under reduced pressure. Purification by prep-HPLC afforded ((1-(6,7-dimethoxyquinazolin-4-yl)piperidin-4-yl)methyl)phosphonic acid **16** (125 mg, 28%) as light yellow solid. LCMS: $[M + H]^+$ *m/z* 368.10. $^1$H NMR (400 MHz, DMSO-$d_6$) δ 8.72 (s, 1H), 7.29 (s, 2H), 4.60 (d, *J* = 12.8 Hz, 2H), 3.95 (d, *J* = 11.2 Hz, 6H), 3.46 (s, 2H), 2.09 (s, 3H), 1.61 (s, 2H), 1.42 (s, 2H).

**Preparation of (2-(1-(6,7-dimethoxyquinazolin-4-yl)piperidin-4-yl)propyl)phosphonic acid 14.**

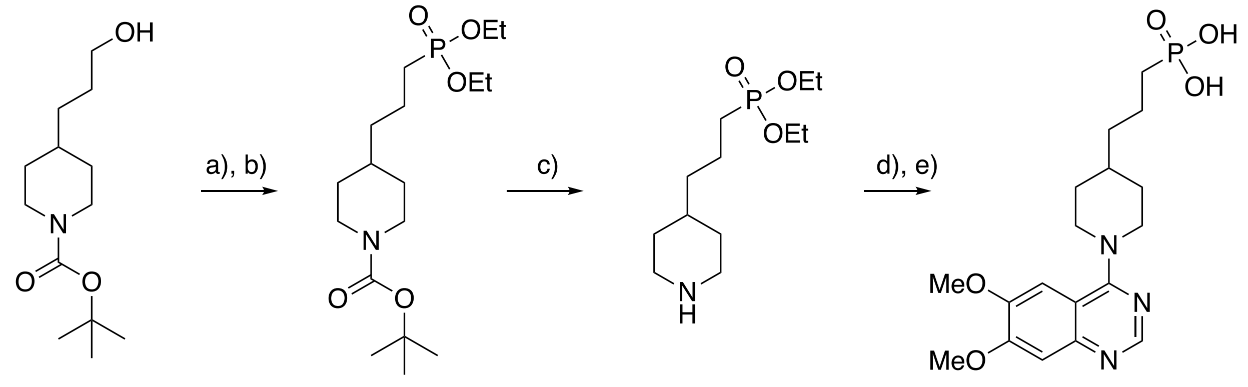


a) $PPh_3$, $I_2$, Imidazole, $CH_2Cl_2$; b)P(O)H(OEt)$_2$, $Cs_2CO_3$, DMF; c) TFA, $CH_2Cl_2$; d) **63**, DIPEA, $CH_2Cl_2$; e) TMSBr, MeCN, 60 °C

Iodine (1.35 g, 5.34 mmol) was added to a solution of $PPh_3$ (1.4 g, 5.34 mmol) and imidazole (0.36 g, 5.34 mmol in $CH_2Cl_2$ (20 mL). The mixture was stirred at rt for 0.5 h and then a solution of **79** (1.0 g, 4.11 mmol) in $CH_2Cl_2$ (5 mL) was added dropwise. The reaction mixture was stirred at rt for 4h and then quenched with saturated $Na_2SO_3$ solution, and extracted with $CH_2Cl_2$. The organic phase was washed with water, brine, dried ($Na_2SO_4$) and evaporated to dryness under reduced pressure. Chromatography ($SiO_2$; 5% EtOAc in petroleum ether) to afford *tert*-butyl 4-(3-iodopropyl)piperidine-1-carboxylate (1.0 g, 68% yield) as light yellow oil.

To a mixture of *tert*-butyl 4-(3-iodopropyl)piperidine-1-carboxylate (1.0 g, 2.83 mmol in DMF (50 mL) was added diethyl phosphonate (0.58 g, 4.24 mmol) and $Cs_2CO_3$ (1.84 g, 5.66 mmol). The reaction mixture was stirred under an atmosphere of nitrogen at rt overnight and then quenched by the addition or water. The organic phase was washed with water, brine, dried ($Na_2SO_4$) and evaporated to dryness under reduced pressure. Chromatography ($SiO_2$; 20% EtOAc in petroleum ether) gave **80** (0.78 g, 76%) as light yellow oil. LCMS: $[M + H]^+$ *m/z* 364.30.

To a solution of **80** (0.78 g, 2.14 mmol) in $CH_2Cl_2$ (8 mL) was added TFA (1.5 mL, 21.4 mmol). The mixture was stirred at rt for 4h and then evaporated to dryness under reduced pressure. To give crude **81** as an oil which was used in the next step without further purification. LCMS: [M + H]$^+$ *m/z* 264.25.

DIPEA (1.37 g, 10.63 mmol) was added to a solution of diethylphosphonate (597 mg, 2.65 mmol) and crude **81** in $CH_2Cl_2$ (10 mL). The mixture was stirred at rt overnight and then quenched with sat'd aqueous $NH_4Cl$ solution and extracted with $CH_2Cl_2$. The organic phase was washed with water, brine, dried ($Na_2SO_4$) and evaporated to dryness under reduced pressure. Chromatography ($SiO_2$, 5% MeOH in $CH_2Cl_2$) gave the diethylphosphonate (0.5 g, 39%) intermediate as a yellow oil. This was solvated in MeCN (10 mL) was TMSBr (1.46 mL, 11.07 mmol) was added. The resulting mixture was stirred at 60 ºC for 6 h and then cooled to room temperature and evaporated to dryness under reduced pressure. Chromatography (Prep-HPLC) gave **14** (220 mg, 50%) as white solid. LCMS: [M + H]$^+$ *m/z* 396.20. $^1$H NMR (400 MHz, Methanol-$d_4$) δ 8.51 (s, 1H), 7.33 (s, 1H), 7.14 (s, 1H), 4.02 (s, 3H), 3.97 (s, 3H), 3.49 (t, *J* = 12 Hz, 2H), 2.00–1.97 (m, 3H), 1.75–1.66 (m, 5H) and 1.45–1.37 (m, 5H).

**Preparation of (2-(1-(6,7-dimethoxyquinazolin-4-yl)piperidin-4-yl)ethyl)phosphonothioic *O*,*O*-acid 17.**

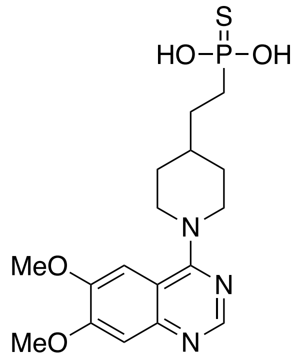

To a stirred solution of *O,O*-diethyl (2-(piperidin-4-yl)ethyl)phosphonothioate (400 mg, 1.51 mmol) and DIPEA (927 mg, 7.19 mmol) in DMSO (10 mL) was added 4-chloro-6,7-dimethoxy-quinazoline **66** (403 mg, 1.80 mmol). The reaction mixture was placed under a nitrogen atmosphere and then stirred at 80 °C for 16 h. The reaction mixture was cooled to room temperature, diluted with water and extracted with ethyl acetate. The organic phase was dried ($Na_2SO_4$) and evaporated to dryness under reduced pressure. Purification ($SiO_2$, 0–100% EtOAc in Hexanes) provided *O,O*-diethyl(2-(1-(6,7-dimethoxyquinazolin-4-yl)piperidin-4-yl)ethyl)phosphonothioate (380 mg, 46%). A stirred solution of *O,O*-diethyl(2-(1-(6,7-dimethoxyquinazolin-4-yl)piperidin-4-yl)ethyl)phosphonothioate (45 mg, 0.099 mmol) in TMSI (7 mL) was stirred at 60 °C for 16 h and then cooled to room temperature. The mixture was diluted with water and extracted with ethyl acetate. The organic phase was dried ($Na_2SO_4$) and then evaporated to dryness under reduced pressure. Purification by prep-HPLC afforded (2-(1-(6,7-dimethoxyquinazolin-4-yl)piperidin-4-yl)ethyl)phosphonothioic *O,O*-acid **17** (13 mg, 32%) as a white solid. LCMS: $[M + H]^+$ *m/z* 396.25. $^1H$ NMR (400 MHz, DMSO-$d_6$) δ 8.51 (s, 1H), 7.33 (s, 1H), 7.16 (s, 1H), 4.02 (s, 3H), 3.97 (s, 3H), 3.58 (d, $J$ = 10.4 Hz, 3H), 3.48 (t, $J$ = 12.0 Hz, 2H), 2.00 (d, $J$ = 11.7 Hz, 2H), 1.81 (s, 1H), 1.64 (d, $J$ = 17.9 Hz, 2H), 1.61–1.51 (m, 2H), 1.45–1.32 (m, 2H).

**(2-(4-(6,7-Dimethoxyquinazolin-4-yl)piperidin-1-yl)ethyl)phosphonic acid 18.**

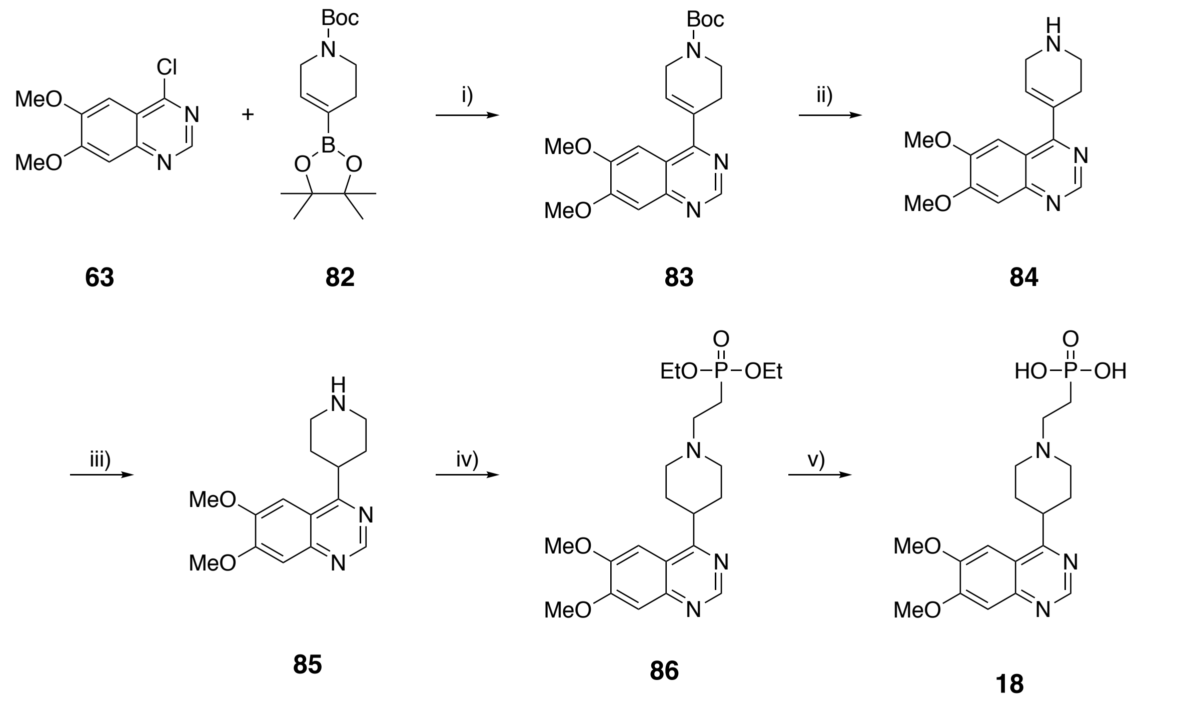


i) $PdCl_2$(dppf), $K_2CO_3$, 1,4-dioxane/$H_2O$; ii) TFA, $CH_2Cl_2$; iii) Pd/C, $H_2$, MeOH; iv) $Na_2CO_3$, $H_2O$, Δ; v) TMSBr, $Na_2CO_3$, $CH_2Cl_2$.

4-Chloro-6,7-dimethoxyquinazoline **63** (1.68 g, 7.5 mmol), *tert*-butyl 4-(4,4,5,5-tetramethyl-1,3,2-dioxaborolan-2-yl)-3,6-dihydropyridine-1(2H)-carboxylate **82** (3.48 g, 11.25 mmol), $PdCl_2$(dppf)-DCM (306 mg, 0.375 mmol, 5 mol%), and $Na_2CO_3$ (1.59 g, 15.0 mmol) were placed in a round-bottomed flask and evacuated and backfilled with $N_2$. 1,4-Dioxane (degassed, 30 mL) was added via syringe followed by the addition of water (degassed, 3 mL) in a similar manner. The reaction mixture was heated at 80 °C for 6 h, cooled to room temperature and filtered through a thin pad of Celite®. The filtrate was washed with EtOAc, washed with brine, dried $Na_2$SO3) and evaporated to dryness under reduced pressure. Chromatography ($SiO_2$; EtOAc in hexanes 1:1) to afford *tert*-butyl 4-(6,7-dimethoxyquinazolin-4-yl)-3,6-dihydropyridine-1(2H)-carboxylate **83** (1.35 g 48%) as a solid. $^1$H NMR (400 MHz, $CDCl_3$) δ 9.07 (s, 1H), 7.36 (s, 1H), 7.33 (s 1H), 6.17 (s, 1H), 4.20 (s, 1H), 4.06 (s, 1H), 3.40 (s, 1H), 3.74 (t, 2H), 2.76–2.73 (m, 2H), 1.51 (s, 9H).

Trifluoroacetic acid (20 mL) was added dropwsie to a mixture of *tert*-butyl 4-(6,7-dimethoxyquinazolin-4-yl)-3,6-dihydropyridine-1(2H)-carboxylate **83** (2.7 g) in dichloromethane (30 mL) at 0 °C. The mixture was stirred at room temperature for 4 h and then diluted with dichloromethane and washed with water, dried ($Na_2SO_3$) and evaporated to dryness under reduced pressure to afford 6,7-dimethoxy-4-(1,2,3,6-tetrahydropyridin-4-yl)quinazoline **84** (3.0 g), which was used for next step without further purification.

$^1$H NMR (400 MHz, $CDCl_3$) δ 9.20 (s, 1H), 8.99 (s, 1H), 7.70 (s 1H), 7.52 (s, 1H), 6.40 (s, 1H), 4.21 (s, 3H), 4.06 (s, 3H), 3.74 (s, 2H), 3.11 (s, 2H), 1.34–1.25 (m, 2H).

A mixture of 6,7-dimethoxy-4-(1,2,3,6-tetrahydropyridin-4-yl)quinazoline **84** (3.0 g, crude), and 10% Pd/C (0.3 g) in MeOH (30 mL) was placed under an atmosphere of hydrogen and stirred at room temperature for 6 hrs. The Pd/C was filtered off through a thin pad of Celite®, washing with MeOH and the filtrate was evaporated to dryness under reduced pressure to afford 6,7-dimethoxy-4-(piperidin-4-yl)quinazoline **85** (2.6 g, 87%) as an off-white solid. $^1$H NMR (400 MHz, $CDCl_3$) δ 9.03 (s, 1H), 7.57 (s, 1H), 7.34 (s 1H), 4.05 (s, 3H), 4.03 (s, 3H), 4.06 (s, 1H), 3.58–3.53 (m, 2H), 3.34–3.33 (d, $J$ = 4.0 Hz, 1H), 3.33–3.30 (d, $J$ = 4.0 Hz, 1H), 2.30–2.12 (m, 5H).

To a mixture of 6,7-dimethoxy-4-(piperidin-4-yl)quinazoline **85** (2.0 g, 5.16 mmol) and diethyl vinylphosphonate (1.3 g, 5.67 mmol) in water (20 mL) was added aqueous $Na_2CO_3$ solution until pH>7 was observed. The mixture was then heated to reflux overnight. The cooled reaction micture was diluted with EtOAc, washed with water, brine and dried ($Na_2SO_3$) and then evaporated to dryness under reduced pressure. Chromatography ($SiO_2$; 10 to 100% EtOAc in Hexanes) provided diethyl (2-(4-(6,7-dimethoxyquinazolin-4-yl)piperidin-1-yl)ethyl)phosphonate **86** (0.50 g, 22%).

To a solution of diethyl (2-(4-(6,7-dimethoxyquinazolin-4-yl)piperidin-1-yl)ethyl) phosphonate **86** (0.41 g, 9.38 mmol) in dichloromethane (30 mL) at 0 °C was added bromotrimethylsilane (14.3

g, 93.8 mmol) dropwise. The reaction was allowed to warm to room temperature and stirred for 6h and then concentrated under reduced pressure. The residue was adjusted to pH>7 with saturated aqueous $Na_2CO_3$ solution. Purification (prep-HPLC) provided the sodium salt of **18** (0.35 g, 92%) as a white solid.LCMS: $[M + H]^+$ *m/z* 382.25 $^1H$ NMR (400 MHz, $D_2O$) δ 8.65 (s, 1H), 6.93 (s, 1H), 6.76 (s, 1H), 3.86 (s, 3H), 3.84 (s, 3H), 3.30–3.22 (m, 1H), 3.18–3.15 (m, 2H), 2.73–2.66 (m, 2H), 2.37–2.32 (m, 2H) and 1.79–1.63 (m, 6H).

**(2-(4-(6,7-Dimethoxyquinazolin-4-yl)piperazin-1-yl)ethyl)phosphonic acid hydrogen bromide salt 19.**

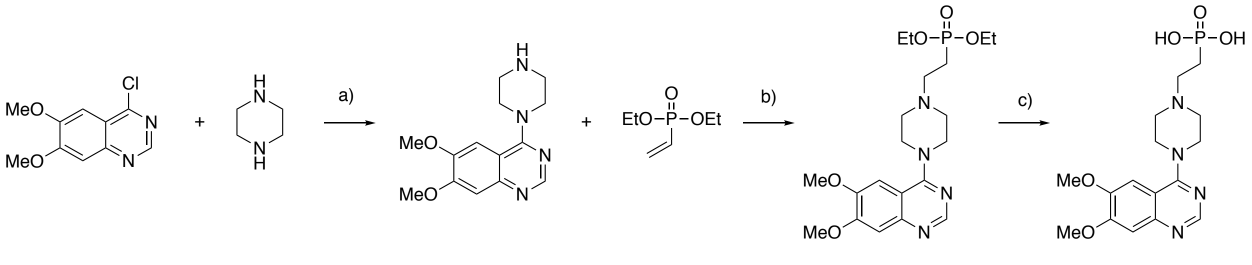


63 87 88 89 90 19

a) iPrOH, Δ; b) $H_2O$, 50 ºC; c) TMSBr, $CHCl_3$, DMF

A mixture containing pyrazine **87** and compound **63** in propan-2-ol were heated at reflux for 30 min and then cooled to room temperature. The reaction mixture was quenched with water and extracted into chloroform. The organic phase was separated, washed with water, brine, dried ($Na_2SO_4$) and evaporated to dryness under reduced pressure. Crystallization from diethyl ether gave **87** as a white solid (1.65 g, 84%). The piperazine **87** (0.31 g, 1.1 mmol) was dissolved in water (20 mL) and vinyl phosphonate **89** (0.19 g, 1.2 mmol) was added. The resulting mixture was heated at 50 ºC for 1 h and then cooled to room temperature. Extraction with chloroform, dried ($Na_2SO_4$) and evaporated to dryness under reduced pressure. Chromatography ($SiO_2$ 12 g; 15% MeOH in $CH_2Cl_2$), followed by crystallization from diethyl ether, gave the ethyl ester **90**

(0.23 g; 49%) as a white solid. LCMS: $[M + H]^+$ *m/z* 410.10. $^1$H NMR (500 MHz, Chloroform-*d*) δ 8.67 (s, 1H), 7.25 (s, 1H), 7.09 (s, 1H), 4.04 (s, 3H), 4.02 (d, *J* = 3.9 Hz, 2H), 4.00 (s, 3H), 3.77 (d, *J* = 10.9 Hz, 6H), 3.68 (t, *J* = 4.9 Hz, 4H), 3.49 (s, 2H), 2.81–2.74 (m, 2H), 2.71 (t, *J* = 4.85 Hz, 4H), 2.11–2.00 (m, 2H).Trimethylsilyl bromide (198 mg, 1.3 mmol) was added to a solution of the 4-[4-(2-diethoxyphosphorylethyl)piperazin-1-yl]-6,7-dimethoxy-quinazoline **90** (600 mg, 3.8 mmol) in chloroform (20 mL) and DMF (5 mL). The resulting solution was left to stir at room temperature for 3 h and then quenched by addition of methanol. The mixture was evaporated to dryness under reduced pressure and crystallized from methanol-diethyl ether to give the desired product **19** (0.23 g, 89%) as the HBr salt. LCMS: $[M + H]^+$ *m/z* 382.8. $^1$H NMR (500 MHz, $D_2O$) δ8.70 (s, 1H), 7.36 (s, 1H), 7.29 (s, 1H), 5.01 (d, *J* = 12.4 Hz, 2H), 4.08 (s, 3H), 4.03 (s, 3H), 4.01–3.96 (m, 2H), 3.88–3.81 (m, 2H), 3.50–3.45 (m, 2H), 2.73 (m, 2H), and 2.19–2.11 (m, 2H).

**Preparation of (4-(6,7-dimethoxyquinazolin-4-yl)phenethyl)phosphonic acid 21.**

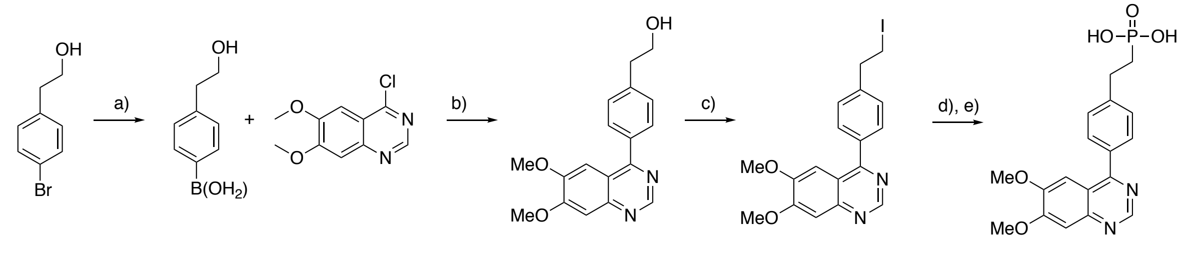


**91** **92** **63** **93** **94** **20**

a) nBuLi, triisopropylborate, THF; b) $Pd(PPh_3)_4$, $K_2CO_3$, $H_2O$, THF, 65 ºC; c) $PPh_3$, imidazole, $I_2$, $CH_2Cl_2$; d) $(BnO)_2P(O)H$, $Cs_2CO_3$, DMF; e) Pd/C, $H_2$, MeOH.

2.5M n-Butyllithium (24 mL) was added to a solution of 2-(4-bromophenyl)ethan-1-ol **91** (5.0 g, 24.8 mmol) in anhydrous THF (100 mL) at –78 ºC under a nitrogen atmosphere. After being stirred for 1 h, triisopropyl borate (8.6 mL) was added to the mixture. The reaction mixture was stirred at room temperature for 1 h and then quenched by the addition of 2M HCl solution (100 mL) and stirred for 1 h. The mixture was extracted with dichloromethane (3 × 100 mL), dried ($Na_2SO_4$) and evaporated to dryness under reduced pressure. Chromatography ($SiO_2$; dichloromethane: methanol, 1:0 to 20:0) gave the boronic acid **92** (1.34 g, 33%) as light yellow solid. This material was then dissolved in a solution of THF (30 mL) and water (10 mL). 4-Chloro-6,7-dimethoxyquinazoline **63** (2.24 g, 10.0 mmol) and potassium carbonate (2.76 g, 20.0 mmol) were added to the solution followed by tetrakis(triphenylphosphine)palladium (0.5 g, 0.43 mmol). The resulting mixture was stirred at 65 ºC for 16 h and then diluted with ethyl acetate, washed with brine, dried ($Na_2SO_4$) and evaporated to dryness under reduced pressure. Chromatography ($SiO_2$: (methanol in dichloromethane 0 to 10%) gave **93** (1.43 g, 60%) as light yellow solid.

To a solution of triphenylphosphine (2.36 g, 9.0 mmol) in dichloromethane (24 mL) was added imidazole (700 mg, 10.28 mmol) at 0 ° C. After being stirred for 10 min, $I_2$ (2.3 g, 9.0 mmol) was added. After being stirred for a further 10 min, compound **93** (1.5 g, 4.8 mmol) in dichloromethane (12 mL). The mixture was allowed to warm to room temperature and stirred for 5 h. Then the mixture was diluted with dichloromethane (36 mL), washed with brine, dried ($Na_2SO_4$) and evaporated to dryness under reduced pressure. Chromatography ($SiO_2$: petroleum ether:ethyl acetate 10:1) gave **94** (4.0 g), as colorless oil.

Cesium carbonate (1.426 g, 4.4 mmol) was added to a mixture of crude **94** (930 mg, 2.2 mmol) and dibenzyl phosphonate (884 mg, 3.37 mmol) in DMF (20 mL). The mixture was placed under an atmosphere of nitrogen and stirred at room temperature for 3 h. Once complete, the reaction mixture was filtered and evaporated to dryness under reduced pressure. Chromatography (C18 column: water:acetonitrile, 1:0 to 80:1) followed by lyophilization gave the dibenzyl intermediate (750 mg, 79%) as off-white solid. Dibenzyl (4-(6,7-dimethoxyquinazolin-4-yl)phenethyl)phosphonate (230 mg, 0.41 mmol) was dissolved in MeOH (20 mL). Pd/C (46 mg,

20% w/w) was added and the mixture stirred under a hydrogen atmosphere at room temperature for 24 h and then filtered through Celite®. Chromatography (prep-HPLC under acidic conditions) gave compound **20** (55.4 mg, 36%) as yellow solid. LCMS: $[M + H]^+$ *m/z* 375.0. $^1H$ NMR (400 MHz, DMSO-$d_6$)): $\delta$ 9.09 (s, 1H), 7.75-7.73 (d, $J$ = 8.0 Hz, 2H), 7.45–7.43 (m, 2H), 7.41 (s, 1H), 7.32 (s, 1H), 4.08 (s, 1H), 3.98 (s, 3H), 3.91 (s, 1H), 3.81 (s, 3H), 2.89–2.87 (m, 3H) and 1.89 (m, 2H).

**Synthesis of (4-((6,7-dimethoxyquinolin-4-yl)amino)phenyl)phosphonic acid sodium salt 21.**

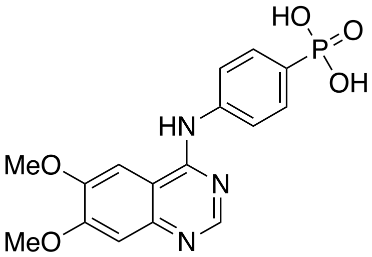


A mixture of 4-chloro-6,7-dimethoxy-quinazoline **63** (0.67 g, 3.0 mmol) and diethyl (4-aminophenyl)phosphonate (0.69 g, 3.0 mmol) in *i*PrOH (10 mL) was heated at reflux overnight. The solid precipitate was filtered, washed with EtOAc and dried to give diethyl (4-((6,7-dimethoxyquinazolin-4-yl)amino)phenyl)phosphonate (0.92 g, 73% yield) as a white solid. The product was dissoved in MeCN (20 mL) and to this was added trimethylsilylbromide (2.8 mL, 22 mmol). The resulting mixture was stirred at 60 °C for 6 h, cooled and then evaporated to dryness under reduced pressure. The crude residue was quenched by the addition of saturated aqueous $NaHCO_3$ (adjusted to pH 8). The resulting mixture was purified by prep-HPLC (under neutral conditions) and then lyophilized to give the desired product **21** (300 mg, 35% yield) as an off-white solid. LCMS: $[M + H]^+$ *m/z* 362.10. $^1H$ NMR (500 MHz, $D_2O$) δ7.96 (s, 1H), 7.70 (dd, $J$ =

9.1, 6.8 Hz, 2H), 7.41 (dd, $J$ = 6.8, 1.7 Hz, 2H), 6.70 (s, 1H), 6.45 (s, 1H), 3.63 (s, 3H), and 3.61 (s, 3H).

**Synthesis of (4-((6,7-dimethoxyquinolin-4-yl)amino)benzyl)phosphonic acid sodium salt 22.**

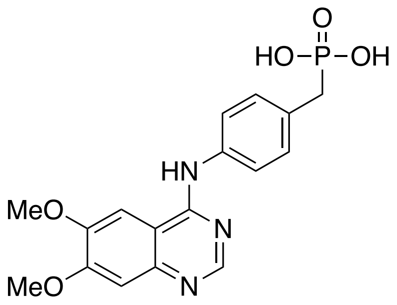


A mixture of 4-chloro-6,7-dimethoxy-quinazoline **63** (0.34 g, 1.5 mmol) and diethyl (4-aminobenzyl)phosphonate (0.36 g, 3.0 mmol) in iPrOH (10 mL) was heated at reflux overnight. The precipitate was filtered, washed with EtOAc and evaporated to dryness under reduced pressure and then dissolved in acetonitrile (20 mL). To this was added trimethylsilylbromide (0.58 mL, 4.6 mmol). The mixture was stirred at 60 °C for 6 h. After concentration, the residue was treated with sat'd aqueous $NaHCO_3$ until the solution reached pH 8. The mixture was purified by prep-HPLC (neutral) to give **22** (104 mg, 57%) as an off-white solid. LCMS: $[M + H]^+$ $m/z$ 376.10. $^1H$ NMR (500 MHz, $D_2O$) δ8.03 (s, 1H), 7.40–7.36 (m, 4H), 6.80 (s, 1H), 6.52 (s, 1H), 3.72 (s, 6H), and 2.92 (d, J = 15.8 Hz, 2H).

**(4-(((6,7-Dimethoxyquinolin-4-yl)amino)methyl)phenyl)phosphonic acid sodium salt 23.**

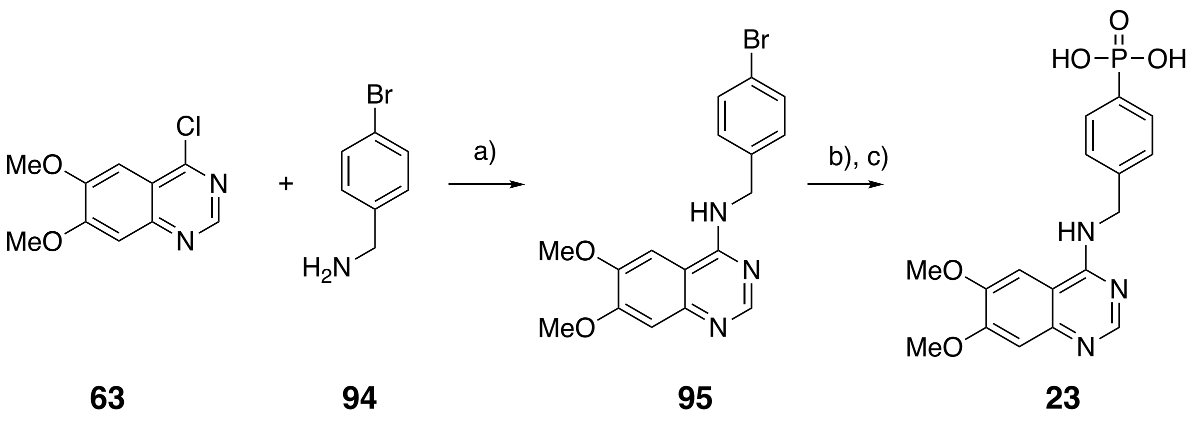

a) iPrOH, Δ; b) $Et_3N$, KOAc, $Pd(OAc)_2$, dppf, $(EtO)_2PH$, THF; c) TMSBr, MeCN, 60 °C.

A mixture of 4-chloro-6,7-dimethoxy-quinazoline **63** (0.93 g, 4.14 mmol) and (4-bromophenyl)methanamine **94** (0.77 g, 4.14 mmol) in iPrOH (10 mL) was heated at reflux overnight. The solid precipitated was filtered; washed with ethyl acetate and evaporated to dryness under reduced pressure to give *N*-(4-bromobenzyl)-6,7-dimethoxyquinazolin-4-amine hydrochloride **95** (1.5 g, 88%) as a white solid. Triethylamine (0.37 mL, 2.68 mmol) was added to a mixture of KOAc (11 mg, 0.112 mmol), $Pd(OAc)_2$ (5.5 mg, 0.025 mmol), dppf (27 mg, 0.049 mmol) in THF (10 mL) was purged with nitrogen. Triethylamine (0.37 mL, 2.68 mmol) was added. After stirring at 70 °C for 15 min, a solution of *N*-(4-bromobenzyl)-6,7-dimethoxyquinazolin-4-amine hydrochloride (0.5 g, 1.22 mmol) and diethyl phosphonate (0.16 g, 1.22 mmol) in THF (10 mL) was added. The reaction was stirred at reflux for 6 h and then partitioned between EtOAc (30 mL) and water (20 mL). The organic phase was separated, washed with water, brine, dried ($Na_2SO_4$) and evaporated to dryness under reduced pressure. Purification by column chromatography ($SiO_2$; 50% petroleum ether in ethyl acetate) afforded diethyl (4-(((6,7-dimethoxyquinazolin-4-yl)amino)methyl)phenyl) phosphonate (0.2 g, 38%) as yellow solid.

To a solution of diethyl (4-(((6,7-dimethoxyquinazolin-4-yl)amino)methyl)phenyl) phosphonate (0.5 g, 1.16 mmol) in MeCN (20 mL) was added TMSBr (1.45 mL, 11.5 mmol). The mixture was stirred at 60 °C for 6 h, cooled to room temperature and then evaporated under reduced pressure. The residue was quenched with saturated aqueous $NaHCO_3$ (pH 9) and the resulting mixture purified prep-HPLC (neutral) to afford the title product **23** as an off-white solid (102 mg, 22%). LCMS: $[M + H]^+$ *m/z*: 376.0 $^1$H NMR (400 MHz, $D_2O$) δ 8.00 (s, 1H), 7.61 (s, 2H), 7.29 (d, *J* = 7.6 Hz, 2H), 6.70 (s, 1H), 6.55 (s, 1H), 4.62 (s, 2H), 3.75 (d, *J* = 18.2 Hz, 6H).

**General procedure synthesis of dimethyl (2-(piperidin-4-yl)ethyl)phosphonate 99 and diethyl (2-(piperidin-4-yl)ethyl)phosphonate 100.**

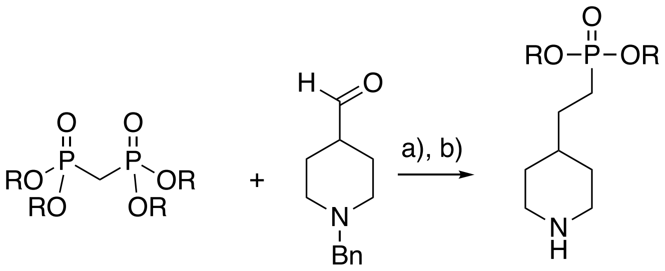


**96** R = Me
**97** R = Et

**98**

**99** R = Me
**100** R = Et

a) NaH, PhMe; b) Pd/C, $H_2$, EtOH.

Sodium hydride (1.1 mol. equiv.) is carefully added to a stirred solution of bis(dimethoxyphosphoryl)methane **96** or bis(diethoxyphosphoryl)methane **97** (1 mol. equiv.) in toluene at room temperature. The reaction mixture is then placed under an atmosphere of nitrogen and a solution of 1-benzylpiperidine-4-carbaldehyde **98** (1 mol. equiv.) in toluene was slowly added, keeping the temperature below 40 ºC. The resulting mixture is left to stir at room temperature for 16 h and then quenched by the addition of aqueous saturated $NH_4Cl$ solution. The organic phase was separated, washed with brine, dried ($MgSO_4$) and evaporated to dryness. Chromatography (120 g $SiO_2$; 5 to 100% gradient of EtOAc in Hexanes) provides dimethyl or diethyl (*E*)-(2-(1-benzylpiperidin-4-yl)vinyl)phosphonates as a colorless oil. To a mixture of dimethyl or diethyl (*E*)-(2-(1-benzylpiperidin-4-yl)vinyl)phosphonate (1 mol. equiv.) in ethanol is added catalytic Pd/C. The mixture is placed under an atmosphere of hydrogen and stirred at room temperature for 12 h, filtered and evaporated to dryness under reduced pressure to give either the dimethyl or diethyl (2-(piperidin-4-yl)ethyl)phosphonates **99** and **100** as colorless oils.

**General procedure for the synthesis of dibenzyl (2-(piperidin-4-yl)ethyl)phosphonate 104.**

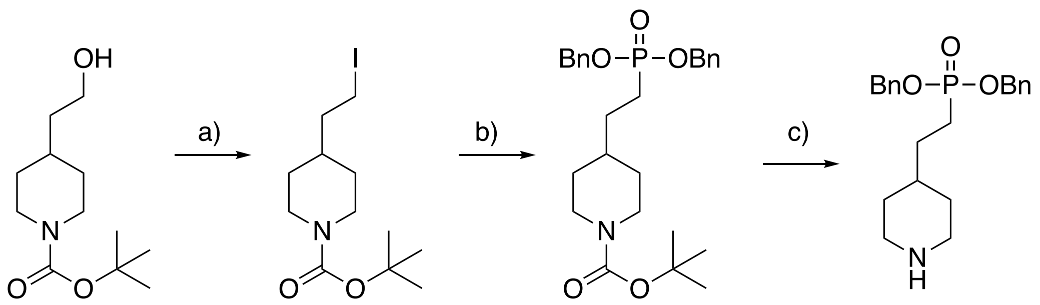


**101** **102** **103** **104**

a) $PPh_3$, $I_2$, imidazole, $CH_2Cl_2$; b) P(O)OBn$_2$, DBU, MeCN; c) TFA, $CH_2Cl_2$

Iodine (1.5 mol. equiv.) was added to a solution of $PPh_3$ (1.5 mol. equiv.) and imidazole (1.5 mol. equiv.) in $CH_2Cl_2$. After stirring for 10 min, a solution of **101** (1.0 mol. equiv.) in $CH_2Cl_2$ is added dropwise. The mixture is stirred at room temperature for 2 h, filtered through a pad of Celite® and treated with 5% sodium thiosulfate solution. The mixture is extracted with ethyl acetate, washed with brine, dried ($Na_2SO_4$) and evaporated to dryness under reduced pressure. Chromatography provides **102** as an oil.

DBU (5.0 mol. equiv.) was added to a solution of compound **102** (3.0 mol. equiv.) in MeCN at 40 ºC. After being stirred for 10 min, a solution of dibenzylphosphonate (1.0 mol equiv.) in

MeCN was added dropwise. After stirring for 2 hours, the reaction mixture is evaporated to dryness under reduced pressure and purified by chromatography to yield **103**.

A solution of compound **103** (1.0 mol. equiv.) in TFA/ DCM is stirred at room temperature for 1 h and then evaporated to dryness under reduced pressure to afford **104** as an oil.

**General methods to synthesize (2-(1-(quinazolin-4-yl)piperidin-4-yl)ethyl)phosphonic acids, (2-(1-(quinolin-4-yl)piperidin-4-yl)ethyl)phosphonic acids and (2-(1-(isoquinolin-1-yl)piperidin-4-yl)ethyl)phosphonic acids.**

**Method A:**

Diisopropylethylamine (2 mol. equiv.) was added to a mixture of either dimethyl (2-(piperidin-4-yl)ethyl)phosphonate **99** or diethyl (2-(piperidin-4-yl)ethyl)phosphonate **100** (1.1 mol. equiv.) and a 4-chloroquinazoline, 4-chloroquinoline or 1-chloroisoquinoline (1 mol. equiv.) in isopropyl alcohol (0.1 M reaction concentration). After stirring at 90 ºC for 3 h, the reaction mixture was cooled and evaporated to dryness. Purification of silica gel (5% MeOH in dichloromethane) provides the dimethyl or diethyl phosphonates. To a cooled (0 °C) solution of the phosphonate (1 mol. equiv.) in chloroform or dichloromethane (0.5 M reaction concentration) was added trimethylsilyl bromide (3 mol. equiv.). The reaction mixture was allowed to warm to room temperature and, after 90 min, was quenched by the addition of methanol. The mixture was evaporated to dryness under reduced pressure and then solvated in methanol. The reaction mixture was concentrated to half volume, filtered to remove precipitate, and then evaporated to dryness. The residue was crystallized with dichloromethane, filtered and dried under reduced pressure to give the desired phosphonic acid as a bromide salt.

**Method B:**

Diisopropylethylamine (3 mol. equiv.) was added to a mixture of either dimethyl (2-(piperidin-4-yl)ethyl)phosphonate **99** or diethyl (2-(piperidin-4-yl)ethyl)phosphonate **100** (1.1 mol. equiv.) and a 4-chloroquinazoline, 4-chloroquinoline or 1-chloroisoquinoline (1 mol. equiv.) in dichloromethane (0.1 M reaction concentration). After stirring at room temperature overnight, the reaction mixture was quenched by the addition of sat'd aqueous $NH_4Cl$ solution. The organic phase was separated and washed with water and brine, dried ($Na_2SO_4$) and evaporated to dryness under reduced pressure. Purification of silica gel (5% MeOH in dichloromethane) provides the dimethyl or diethyl phosphonates. To a cooled (0 °C) solution of the dimethyl or diethyl phosphonates (7 mol. equiv.) in acetonitrile (0.1 M reaction concentration) was added trimethylsilyl bromide (3 mol. equiv.). The reaction mixture was stirred at 60 °C for 6 h, cooled and evaporated to dryness under reduced pressure and the crude residue quenched by the addition of sat'd aqueous $NaHCO_3$ solution (until pH 8~9 was observed). The crude residue purified by prep-HPLC (neutral) to give the phosphonic acid as a sodium salt.

**Method C:**

Diisopropylethylamine (3 mol. equiv.) was added to a mixture of either dibenzyl(2-(piperidin-4-yl)ethyl)phosphonate **104** (1.1 mol. equiv.) and a 4-chloroquinazoline, 4-chloroquinoline or 1-chloroisoquinoline (1 mol. equiv.) in dichloromethane (0.1 M reaction concentration). After stirring at room temperature overnight, the reaction mixture was quenched by the addition of sat'd aqueous $NH_4Cl$ solution. The organic phase was separated and washed with water and brine, dried ($Na_2SO_4$) and evaporated to dryness under reduced pressure. Purification of silica gel (5% MeOH in dichloromethane) provides the dibenzyl phosphonates. A mixture of the dibenzyl phosphonate

(1 mol. equiv.) and Pd/C in MeOH is placed under an atmosphere of hydrogen and stirred at room temperature for 2 h. The mixture is then filtered through Celite® and evaporated to dryness under reduced pressure to give the phosphonic acids.

**Preparation of (2-(1-(6,7-dimethoxyquinazolin-4-yl)piperidin-4-yl)ethyl)phosphonic acid 15.**

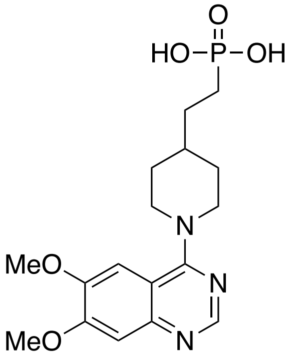


Prepared according to Method A to give **15** (2.1 g, 69%) as an off-white solid. LCMS: $[M + H]^+$ *m/z* 381.8. $^1$H NMR (500 MHz, DMSO-$d_6$) δ 8.77 (s, 1H), 7.34 (s, 1H), 7.23 (s, 1H), 4.71 (d, *J* = 13.1 Hz, 2H), 3.99 (s, 3H), 3.97 (s, 3H), 3.48 (t, *J* = 12.7 Hz, 2H), 3.18 (s, 1H), 1.97–1.90 (m, 2H), 1.62–1.43 (m, 4H), 1.40–1.27 (m, 2H).

**Preparation of (4-(((6,7-dimethoxyquinazolin-4-yl)amino)methyl)benzyl)phosphonic acid 24.**

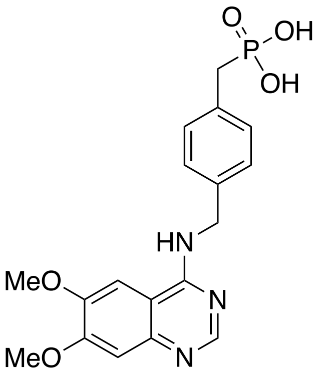

Prepared according to Method B. The product was isolated by prep-HPLC (9% yield) as an off-white solid. LCMS: $[M + H]^+$ *m/z* 390.15. $^1$H NMR (500 MHz, Methanol-$d_4$) δ8.31 (s, 1H), 7.56 (s, 1H), 7.36 (dd, *J* = 8.4, 1.8 Hz, 2H), 7.22 (d, *J* = 8.0 Hz, 2H), 7.08 (s, 1H), 4.74 (s, 2H), 3.96 (s, 3H), 3.93 (s, 3H), and 2.87 (d, *J* = 19.9 Hz, 2H).

**Preparation of (2-(1-(6-methoxyquinazolin-4-yl)piperidin-4-yl)ethyl)phosphonic acid 25.**

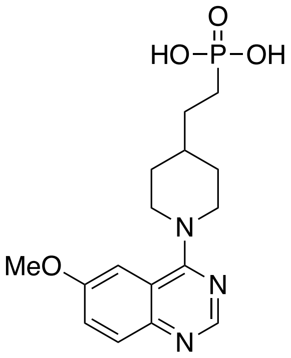

Prepared according to Method A to give **25** (50% yield) as an off-white solid. LCMS: $[M + H]^+$ *m/z* 352.10. $^1$H NMR (400 MHz, Methanol-$d_4$) δ 8.57 (s, 1H), 7.74–7.73 (m, 1H), 7.68–7.66 (m, 1H), 7.46 (d, 1H), 4.96 (br s, 2H), 3.98, (s, 3H), 3.57 (br s, 2H), 2.65 (s, 2H), 2.07–2.04 (m, 2H), 1.81 (m, 1H), 1.79–1.75 (m, 2H), 1.66–1.63 (m, 2H) and 1.46–1.44 (m, 2H).

**Preparation of (2-(1-(6-hydroxyquinazolin-4-yl)piperidin-4-yl)ethyl)phosphonic acid 26.**

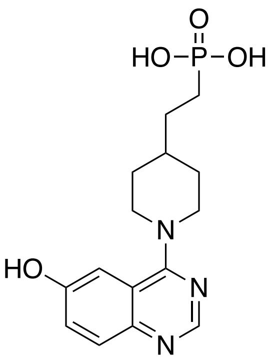

Prepared according to Method C to give **26** (7% yield) as an off-white solid. LCMS: $[M + H]^+$ *m*/z 338.15. $^1$H NMR (400 MHz, DMSO-$d_6$) δ 8.48 (s, 1H), 7.65 (d, *J* = 8.8 Hz, 1H), 7.32 (d, *J* = 8.8 Hz, 1H), 7.18 (s, 1H), 4.21–4.17 (m, 2H), 2.99–2.95 (m, 2H), 1.82–1.79 (m, 2H), 1.53–1.49 (m, 5H) and 1.30–1.19 (m, 2H).

**Preparation of (2-(1-(7-methoxyquinazolin-4-yl)piperidin-4-yl)ethyl)phosphonic acid 27.**

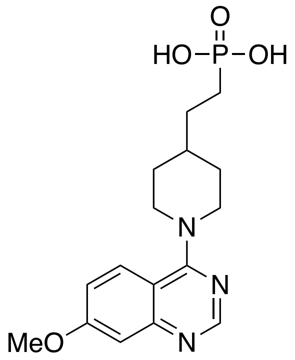

Prepared according to Method A to give **27** (95% yield) as an off-white solid. LCMS: $[M + H]^+$ *m/z* 352.0 $^1$H NMR (500 MHz, Methanol-$d_4$) δ 8.55 (s, 1H), 8.10 (d, *J* = 10 Hz, 1H), 7.30 (dd, *J* = 10 Hz and 5 Hz, 1H), 7.10 (d, *J* = 5 Hz, 1H), 4.01 (s, 3H), 3.57–3.48 (m, 2H), 2.65 (s, 1H), 2.05–2.02 (m, 2H), 1.94–1.90 (m, 1H), 1.80–1.74 (m, 2H), 1.65–1.60 (m, 2H) and 1.46–1.41 (m, 2H).

**Preparation of (2-(1-(7-ethoxyquinazolin-4-yl)piperidin-4-yl)ethyl)phosphonic acid 28.**

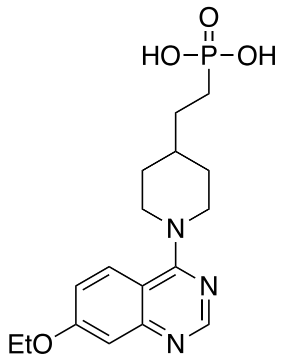

Prepared according to Method B to give **28** (28% yield) as an off-white solid. LCMS: $[M + H]^+$ *m/z* 366.1 $^1H$ NMR (500 MHz, DMSO-$d_6$) δ 8.51 (s, 1H), 7.85 (d, *J* = 7.2 Hz, 1H), 7.14–7.10 (m, 2H), 4.24 (d, *J* = 10.0 Hz, 2H), 4.18 (q, *J* = 5.6 Hz, 2H), 3.07 (t, *J* = 9.6 Hz, 2H), 1.81 (d, *J* = 9.1 Hz, 2H), 1.64–1.44 (m, 5H), 1.34 (t, *J* = 5.6 Hz, 3H), and 1.34–1.25 (m, 2H).

**Preparation of (2-(1-(7-Hydroxyquinazolin-4-yl)piperidin-4-yl)ethyl)phosphonic acid 29.**

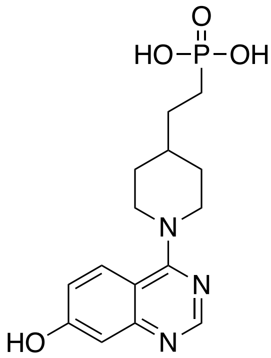


Prepared according to Method B to give **29** (4% yield) as an pale yellow solid. LCMS: $[M + H]^+$ *m/z* 338.25. $^1H$ NMR (400 MHz, DMSO-$d_6$) δ 11.49 (s, 1H), 8.64 (s, 1H), 7.96 (d, *J* = 9.2 Hz, 1H), 7.10 (dd, *J* = 9.2 and 2.2 Hz, 1H), 7.00 (d, J = 2.2 Hz, 1H), 4.62 (br s, 2H), 3.38 (br s, 2H), 1.87 (d, *J* = 12.7 Hz, 2H), 1.72 (br s, 1H), 1.58–1.38 (m, 4H) and 1.28–1.22 (m, 2H).

**Preparation of (2-(1-(7-aminoquinazolin-4-yl)piperidin-4-yl)ethyl)phosphonic acid 30.**

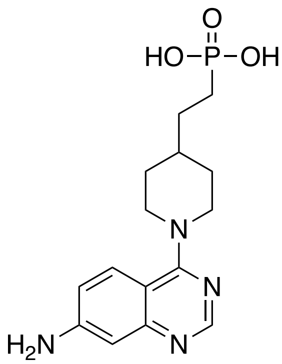

Prepared according to Method C to give **30** (32% yield) as a light yellow solid. LCMS: [M + H]$^+$ *m/z* 337.10. $^1$H NMR (400 MHz, DMSO-$d_6$) δ 8.35 (s, 1H), 7.62 (d, *J* = 8.8 Hz, 1H), 6.81 (d, *J* = 8.8 Hz, 1H), 6.61 (br s, 1H), 6.30 (br s, 2H), 4.26–4.20 (m, 2H), 3.10–2.90 (m, 2H), 1.79–1.76 (m, 2H), 1.60–1.30 (m, 5H) and 1.25–1.20 (m, 2H).

**Preparation of (2-(1-(7-isopropoxyquinazolin-4-yl)piperidin-4-yl)ethyl)phosphonic acid 31.**

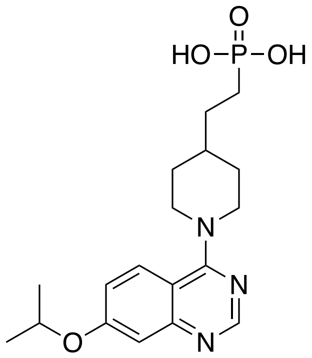


Prepared according to Method B to give **31** (31% yield) as a light-yellow solid. LCMS: [M + H]$^+$ *m/z* 337.10. $^1$H NMR (400 MHz, DMSO-$d_6$) δ 8.35 (s, 1H), 7.62 (d, *J* = 8.8 Hz, 1H), 6.81 (d, *J* = 9.2 Hz, 1H), 6.66 (s, 1H), 6.30 (br s, 2H), 4.25–4.21 (m, 2H), 3.08–2.96 (m, 2H), 1.81–1.75 (m, 2H), 1.65–1.31 (m, 5H) and 1.27–1.18 (m, 2H).

**Preparation of (2-(1-(8-methoxyquinazolin-4-yl)piperidin-4-yl)ethyl)phosphonic acid 32.**

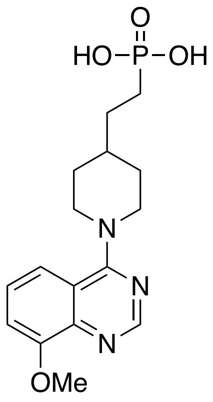

Prepared according to Method B to give **32** (32% yield) as an off-white solid. LCMS: $[M + H]^+$ $m/z$ 352.15. $^1H$ NMR (400 MHz, DMSO-$d_6$) δ 7.92 (s, 1H), 6.92–6.88 (m, 1H), 6.80 (d, $J$ = 7.6 Hz, 1H), 6.71 (d, $J$ = 8.4 Hz, 1H), 3.76–3.70 (m, 2H), 3.71 (s, 3H), 2.72 (t, $J$ = 12 Hz, 2H), 1.64 (d, $J$ = 12 Hz, 2H), 1.51–1.28 (m, 5H) and 1.02–0.94 (m, 2H).

**Preparation of (2-(1-(8-ethoxyquinazolin-4-yl)piperidin-4-yl)ethyl)phosphonic acid 33.**

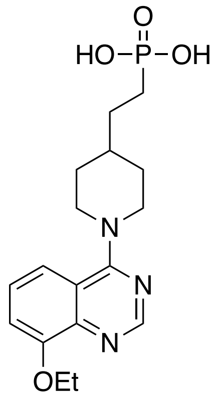


Prepared according to Method B to give **33** as a white solid. LCMS: $[M + H]^+$ $m/z$ 366.20. $^1H$ NMR (400 MHz, DMSO-$d_6$) $^1H$ NMR (500 MHz, $D_2O$) δ8.25 (s, 1H), 7.27–7.21 (m, 2H), 7.12 (d, $J$ = 6.0 Hz, 1H), 4.16 (q, $J$ = 5.6 Hz, 2H), 4.10 (d, $J$ = 10.4 Hz, 2H), 3.04 (t, $J$ = 9.7 Hz, 2H), 1.87 (d, $J$ = 9.2 Hz, 2H), 1.60 (m, 1H), 1.54–1.43 (m, 7H), and 1.36–1.29 (m, 2H).

**Preparation of (2-(1-(8-hydroxyquinazolin-4-yl)piperidin-4-yl)ethyl)phosphonic acid 34.**

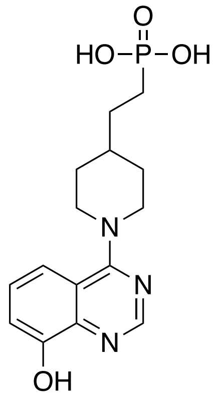

Prepared according to Method B to give **34** (37% yield) as an off-white solid.

LCMS: $[M + H]^+$ *m/z* 338.1 $^1$H NMR (500 MHz, DMSO-$d_6$) δ 8.60 (s, 1H), 7.47–7.42 (m, 2H), 7.27 (d, $J$ = 5.4 Hz, 1H), 4.55 (d, $J$ = 5.4 Hz, 2H), 3.29 (t, $J$ = 9.6 Hz, 2H), 1.88 (d, $J$ = 9.6 Hz, 2H), 1.71 (m, 1H), 1.71–1.57 (m, 4H), and 1.35–1.28 (m, 2H).

**Preparation of (2-(1-(8-isopropoxyquinazolin-4-yl)piperidin-4-yl)ethyl)phosphonic acid 35.**

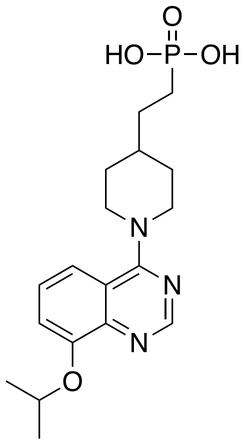


Prepared according to Method B to give **35** (43% yield) as an off-white solid. LCMS: $[M + H]^+$ *m/z* 380.1 $^1$H NMR (400 MHz, DMSO-$d_6$) δ $^1$H NMR (500 MHz, $D_2O$) δ 8.36 (s, 1H), 7.36–7.33 (m, 1H), 7.30–7.26 (m, 2H), 4.83 (sep, $J$ = 4.9 Hz, 1H), 4.16 (d, $J$ = 10.2 Hz, 2H), 3.10 (t, $J$ = 9.7 Hz, 2H), 1.96 (d, $J$ = 9.6 Hz, 2H), 1.67–1.52 (m, 5H), 1.48 (d, $J$ = 4.8 Hz, 6H), and 1.39–1.36 (m, 2H).

**Preparation of (2-(1-(5,8-Dimethoxyquinazolin-4-yl)piperidin-4-yl)ethyl)phosphonic acid 36**.

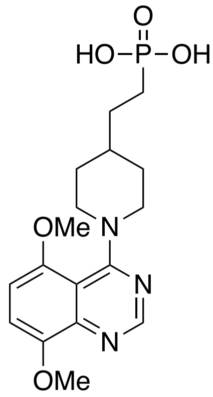

Prepared according to Method B to give **36** as an off-white solid. LCMS: $[M + H]^+$ *m/z* 382.15. $^1$H NMR (500 MHz, $D_2O$) δ8.27 (s, 1H), 7.40 (d, *J* = 7.3 Hz, 1H), 7.03 (d, *J* = 7.3 Hz, 1H), 5.0 (d, *J* = 10.1 Hz, 1H), 3.98 (s, 3H), 3.94 (s, 3H), 3.79 (d, *J* = 10.4 Hz, 1H), 3.42 (d, *J* = 10.2 Hz, 1H), 3.25 (t, *J* = 9.6 Hz, 1H), 2.03–1.97 (m, 2H), 1.87 (d, *J* = 9.5 Hz, 1H), 1.79–1.71 (m, 4H), and 1.58–1.51 (m, 2H).

**Preparation of (2-(1-(6,8-dimethoxyquinazolin-4-yl)piperidin-4-yl)ethyl)phosphonic acid 37.**

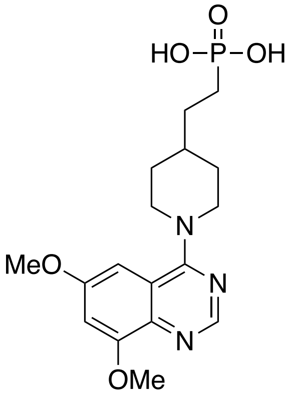

Prepared according to Method C to give **37** (15% yield) as an off-white solid. LCMS: $[M + H]^+$ *m/z* 382.15 $^1$H NMR (500 MHz, DMSO-$d_6$) δ 8.55 (s, 1H), 7.19 (d, *J* = 2.1 Hz, 1H), 6.87 (d, *J* = 2.1 Hz, 1H), 4.65 (d, *J* = 12.7 Hz, 2H), 4.00 (s, 3H), 3.89 (s, 3H), 3.39 (t, *J* = 12.2 Hz, 2H), 1.88 (d, *J* = 12.7 Hz, 2H), 1.72 (m, 1H), 1.55–1.41 (m, 4H), and 1.32–1.25 (m, 2H).

**Preparation of (2-(1-(7,8-dimethoxyquinazolin-4-yl)piperidin-4-yl)ethyl)phosphonic acid 38.**

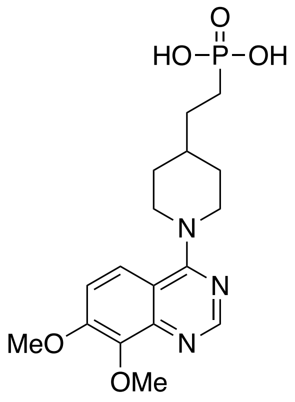

Prepared according to Method C to give **38** (16% yield) as an off-white solid. LCMS: $[M + H]^+$ *m/z* 382.15 $^1$H NMR (400 MHz, DMSO-$d_6$) δ8.60 (s, 1H), 7.90 (d, *J* = 9.6 Hz, 1H), 7.49 (d, *J* = 9.2 Hz, 1H), 4.69 (m, 2H), 4.02 (s, 3H), 3.89 (s, 3H), 3.46 (m, 2H), 1.90 (d, J = 12.8 Hz, 2H), 1.75 (m, 1H), 1.53–1.49 (m, 4 H) and 1.31–1.28 (m, 2H).

**Preparation of (2-(1-(7,8-dihydro-[1,4]dioxino[2,3-g]quinazolin-4-yl)piperidin-4-yl)ethyl)phosphonic acid 39.**

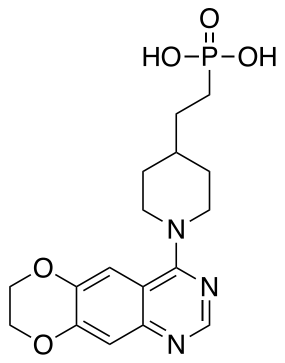

Prepared according to Method C to give **39** (7% yield) as an off-white solid. LCMS: $[M + H]^+$ *m/z* 380.15. $^1$H NMR (400 MHz, DMSO-$d_6$) δ8.64 (s, 1H), 7.48 (s, 1H), 7.19 (s, 1H), 4.56 (d, *J* = 11.8 Hz, 2H), 4.45 (d, *J* = 3.0 Hz, 2H), 4.38 (d, *J* = 3.3 Hz, 2H), 3.39 (s, 1H), 3.33 (s, 1H), 1.87 (d, *J* = 12.2 Hz, 2H), 1.71 (s, 1H), 1.58–1.38 (m, 4H), 1.26 (d, *J* = 10.2 Hz, 2H).

**Preparation of (2-(1-(5-fluoro-8-methoxyquinazolin-4-yl)piperidin-4-yl)ethyl)phosphonic acid 40.**

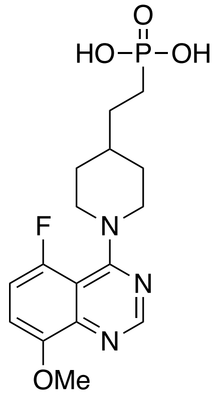

Prepared according to Method C. to give **40** (7% yield) as a light-yellow solid. LCMS: $[M + H]^+$ $m/z$ 370.10. $^1H$ NMR (400 MHz, DMSO-$d_6$) δ8.55 (s, 1H), 7.44–7.38 (m, 2H), 4.28–4.23 (m, 2H), 3.94 (s, 3H), 3.28–3.18 (m, 2H), 1.82–1.78 (m, 2H), 1.70–1.66 (m, 1H), 1.49–1.23 (m, 4H) and 1.26–1.09 (m, 2H).

**Preparation of (2-(1-(6-fluoro-8-methoxyquinazolin-4-yl)piperidin-4-yl)ethyl)phosphonic acid 41.**

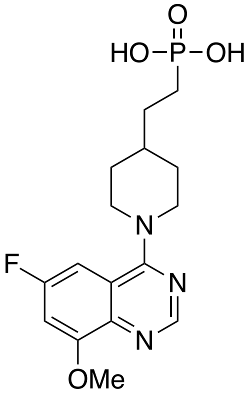

Prepared according to method B to give **41** (44% yield) as a white solid. LCMS: $[M + H]^+$ *m/z* 370.0. $^1$H NMR (500 MHz, $D_2O$) δ7.87 (s, 1H), 6.58 (dd, *J* = 1.7, 8.9 Hz, 1H), 6.37 (dd, *J* = 1.7, 8.2 Hz, 1H), 3.7 (d, *J* = 10.2 Hz, 2H), 3.62 (s, 3H), 2.69 (t, *J* = 9.6 Hz, 2H), 1.58 (d, *J* = 9.4 Hz, 2H), 1.35–1.25 (m, 5H), and 1.03–0.96 (m, 2H).

**Preparation of (2-(1-(6-chloro-8-methoxyquinazolin-4-yl)piperidin-4-yl)ethyl)phosphonic acid 42.**

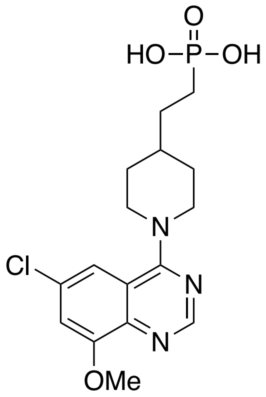


Prepared according to Method B to give **42** (42% yield) as a light-yellow solid. LCMS: $[M + H]^+$ *m/z* 386.10. $^1$H NMR (400 MHz, $D_2O$) δ8.03 (s, 1H), 6.91–6.88 (m, 2H), 3.92–3.89 (m, 2H), 3.77 (s, 3H), 2.93–2.87 (m, 2H), 1.70–1.67 (m, 2H) and 1.41–1.12 (m, 7H).

**Preparation of (2-(1-(7-chloro-8-methoxyquinazolin-4-yl)piperidin-4-yl)ethyl)phosphonic acid 43.**

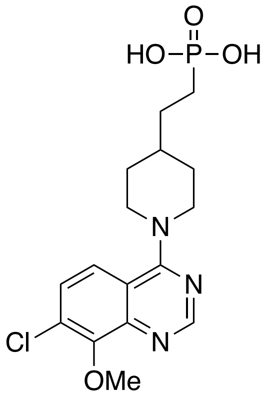

Prepared according to Method B to give **43** (50% yield) as a white solid. LCMS: $[M + H]^+$ *m/z* 386.05. $^1H$ NMR (400 MHz, $D_2O$) δ8.07 (s, 1H), 7.10–7.06 (m, 2H), 3.95–3.91 (m, 2H), 3.71 (s, 3H), 2.96–2.90 (m, 2H), 1.71–1.68 (m, 2H) and 1.42–1.01 (m, 7H).

**Preparation of 2-[1-[6,7-dimethoxy-2-[(*E*)-2-(3-pyridyl)vinyl]quinazolin-4-yl]-4-piperidyl]ethyl-hydroxy-phosphinate 44.**

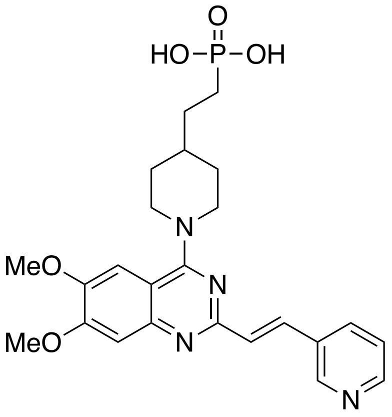

Prepared according to Method B to give **44** (49% yield) as a light-yellow solid. LCMS: $[M + H]^+$ *m/z* 485.25. $^1H$ NMR (400 MHz, $D_2O$) δ8.09 (s, 1H), 7.94 (s, 1H), 7.36 (d, *J* = 8 Hz, 1H), 7.06 (s, 1H), 6.74 (d, *J* = 16.8 Hz 1H), 6.58 (d, *J* = 3.2 Hz, 1H), 6.40 (d, *J* = 3.2 Hz, 1H), 6.24 (d,

$J$ = 16.8 Hz, 1H), 3.94–3.91 (m, 2H), 3.84 (s, 3H), 3.67 (s, 3H), 2.96–2.90 (m, 2H), 1.96-1.93 (m, 2H) and 1.56–1.32 (m, 7H).

**Preparation of (*E*)-(2-(1-(8-Methoxy-2-(2-(pyridin-3-yl)vinyl)quinazolin-4-yl)piperidin-4-yl)ethyl)phosphonic acid 45.**

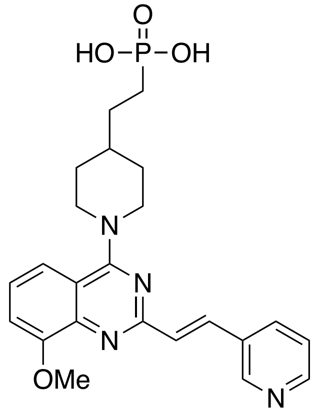


Prepared according to Method B to give **45** (79% yield) as a yellow solid. LCMS: $[M + H]^+$ $m/z$ 455.20. $^1H$ NMR (500 MHz, DMSO-$d_6$) δ9.18 (s, 1H), 8.87 (d, $J$ = 4 Hz, 1H), 8.56 (d, $J$ = 6.5 Hz, 1H), 8.30 (d, $J$ = 13.0 Hz, 1H), 7.95 (m, 1H), 7.80 (d, $J$ = 12.7 Hz, 1H), 7.69–7.63 (m, 3H), 5.01–4.69 (br s, 2H), 4.11 (s, 3H), 3.62–3.44 (br s, 2H), 1.98 (d, $J$ = 9.4 Hz, 2H), 1.87–1.78 (m, 1H), 1.62–1.47 (m, 4H), and 1.44–1.37 (m, 2H).

**Preparation of (2-(1-(6,7-dimethoxyquinolin-4-yl)piperidin-4-yl)ethyl)phosphonic acid 46.**

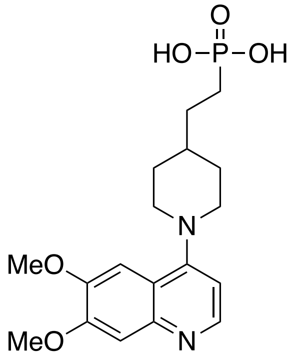

Prepared according to Method C to give **46** (22% yield) as a white solid. LCMS: $[M + H]^+$ *m/z* 381.30. $^1$H NMR (400 MHz, Methanol-$d_4$) δ8.35 (d, $J$ = 6.8 Hz, 1H), 7.29–7.27 (m, 2H), 7.11 (d, $J$ = 6.8 Hz, 1H), 4.27–4.23 (m, 2H), 4.03 (s, 3H), 4.02 (s, 3H), 3.40–3.32 (m, 2H), 2.06–2.03 (m, 4H) 1.82–1.79 (m, 3H) and 1.62–1.48 (m, 2H).

**Preparation of (2-(1-(3-cyano-6,7-dimethoxyquinolin-4-yl)piperidin-4-yl)ethyl)phosphonic acid 47.**

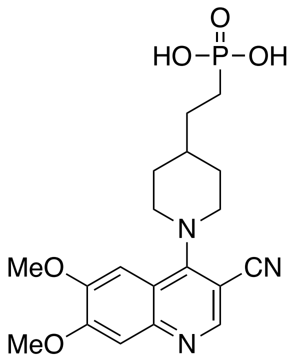


Prepared according to Method B to give **47** (47% yield) as an off-white solid. LCMS: $[M + H]^+$ *m/z* 406.20. $^1$H NMR (400 MHz, $D_2O$) δ8.00 (s, 1H), 6.62 (s, 1H), 6.40 (s, 1H), 3.75 (s, 3H), 3.66 (s, 3H), 3.18 (d, $J$ = 12.3 Hz, 2H), 2.94 (t, $J$ = 12.2 Hz, 2H), 1.72 (d, $J$ = 12.7 Hz, 2H), 1.43–1.30 (m, 6H), 1.17–1.04 (m, 2H).

**Preparation of (2-(1-(3-cyano-6-methoxyquinolin-4-yl)piperidin-4-yl)ethyl)phosphonic acid 48.**

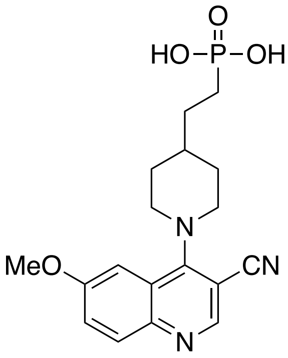

Prepared according to Method B to give **48** (16% yield) as an off-white solid. LCMS: $[M + H]^+$ $m/z$ 376.20. $^1H$ NMR (400 MHz, $D_2O$) δ8.15 (s, 1H), 7.51 (s, 1H), 7.18 (s, 1H), 6.88 (s, 1H), 3.71 (s, 3H), 3.60–3.51 (m, 2H), 3.15–3.08 (m, 2H), 1.81–1.74 (m, 2H) and 1.41–1.15 (m, 7H).

**Preparation of (2-(1-(3-cyano-7-methoxyquinolin-4-yl)piperidin-4-yl)ethyl)phosphonic acid 49.**

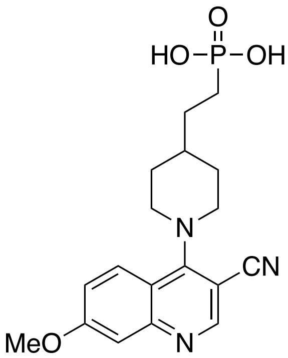

Prepared according to Method B to give **49** (23% yield) as an off-white solid. LCMS: $[M + H]^+$ $m/z$ 376.20. $^1H$ NMR (400 MHz, $D_2O$) δ7.94 (s, 1H), 7.39 (d, $J$ = 9.4 Hz, 1H), 6.84–6.64 (m, 2H), 3.90 (s, 3H), 3.59 (d, $J$ = 12.4 Hz, 2H), 3.22 (t, $J$ = 12 Hz, 2H), 1.89 (d, $J$ = 12.8 Hz, 2H), 1.62–1.45 (m, 5H) and 1.33–1.25 (m, 2H).

**Preparation of (2-(1-(3-cyano-8-methoxyquinolin-4-yl)piperidin-4-yl)ethyl)phosphonic acid 50.**

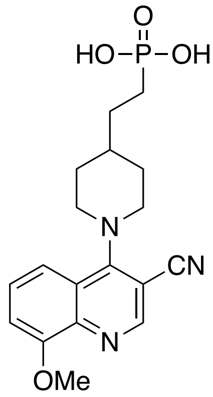


Prepared according to Method B to give **50** (35% yield) as an off-white solid. LCMS: $[M + H]^+$ *m/z* 376.20. $^1$H NMR (400 MHz, $D_2O$) δ8.03 (s, 1H), 7.27–7.23 (m, 1H), 7.11–7.09 (m, 1H), 7.04–7.02 (m, 1H), 3.90 (s, 3H), 3.43 (br d, *J* = 12.4 Hz, 2H), 3.06 (br t, *J* = 12 Hz, 2H), 1.80 (br d, *J* = 12.8 Hz, 2H), 1.50–1.47 (m, 5H) and 1.31–1.24 (m, 2H).

**Preparation of (2-(1-(6,7-dimethoxyisoquinolin-1-yl)piperidin-4-yl)ethyl)phosphonic acid 51.**

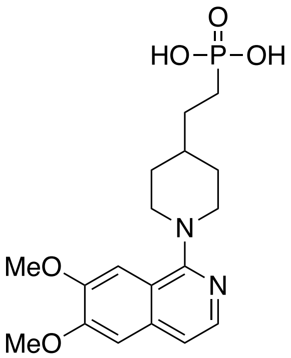


Prepared according to Method B to give **51** (30% yield) as an off-white solid. LCMS: $[M + H]^+$ *m/z* 381.10. $^1$H NMR (400 MHz, $D_2O$) δ $^1$H NMR (400 MHz, DMSO-$d_6$): *δ* 7.79 (d, *J = 6.4 Hz*, 1H), 7.51–7.44 (m, 2H), 7.31 (s, 1H), 3.96 (d, *J* = 4.8 Hz, 6H), 3.23 (m, 4H), 1.88 (m, 2H), 1.55 (m, 2H), 1.48 (m, 1H) and 1.45 (m, 4 H).

**Preparation of (2-(1-(4-cyano-6,7-dimethoxyisoquinolin-1-yl)piperidin-4-yl)ethyl)phosphonic acid 52.**

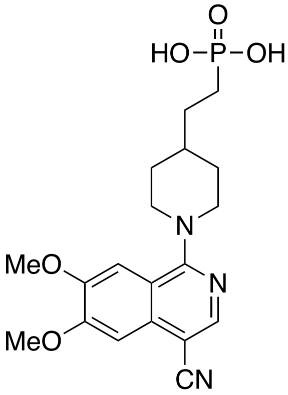

Prepared according to Method B to give **52** (50% yield) as an off-white solid. LCMS: $[M + H]^+$ $m/z$ 406.0 $^1$H NMR (500 MHz, DMSO-$d_6$) δ 8.45 (s, 1H), 7.26 (s, 1H), 7.16 (s, 1H), 4.01 (d, $J$ = 10.6 Hz, 2H), 3.99 (s, 3H), 3.94 (s, 3H), 3.00 (t, $J$ = 9.5 Hz, 2H), 1.83 (d, $J$ = 9.2 Hz, 2H), 1.60–1.48 (m, 5H), and 1.41–1.35 (m, 2H).

**Preparation of *O*-((1-(8-methoxyquinazolin-4-yl)piperidin-4-yl)methyl)*O*,*O*-dihydrogen phosphorothioate 53.**

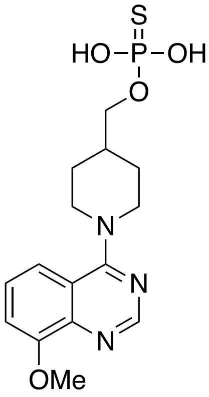

To a solution of (1-(8-methoxyquinazolin-4-yl)piperidin-4-yl)methanol (prepared in using the same method as compound **80**) (500 mg, 1.83 mmol) in pyridine (5 mL) was added phosphorothioyl trichloride (1.6 g, 9.45 mmol) dropwise at –15 ºC. After being stirred at 0 ºC for 1 h, the reaction mixture was added to a solution of sodium bicarbonate (923 mg, 10.98 mmol) in water (20 mL) at 0 ºC. The resulting mixture was stirred at 0 ºC for 2 h. and then evaporated to dryness under reduced pressure. Purification (prep-HPLC) gave **53** (83 mg, in 12%)as a white solid. LCMS: $[M + H]^+$ *m/z* 354.10 $^1H$ NMR (400 MHz, , DMSO-$d_6$) δ8.59 (s, 1H), 7.59–7.50 (m, 2H), 7.45 (dd, *J* = 6.5, 2.4 Hz, 1H), 4.53 (d, *J* = 12.7 Hz, 2H), 3.97 (s, 3H), 3.80–3.74 (m, 4H), 2.03 (s, 1H), 1.86 (d, *J* = 13.5 Hz, 2H), 1.41 (q, *J* = 11.8 Hz, 2H).

**Preparation of (((1-(8-methoxyquinazolin-4-yl)piperidin-4-yl)oxy)methyl)phosphonic acid 54.**

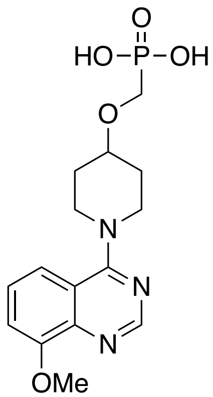


Prepared using the same method as compounds **10** and **11**. The mixture was purified (prep-HPLC 0.1% TFA) to give **54** as an off-white solid.

LCMS: $[M + H]^+$ *m/z* 353.3. $^1H$ NMR (400 MHz, $D_2O$) δ 8.40 (s, 1H), 7.54 (s, 2H), 7.40 (d, *J* = 7.0 Hz, 1H), 4.37 (s, 3H), 4.01–3.88 (m, 9H), 3.71 (d, *J* = 9.3 Hz, 4H), 3.63 (s, 1H), 2.11 (s, 2H), 1.81 (s, 2H).

**Preparation of (4-(8-methoxyquinazolin-4-yl)phenethyl)phosphonic acid 55.**

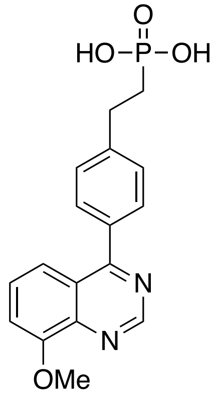

Prepared using the same method as compound **20** as a white solid. LCMS: $[M + H]^+$ *m/z* 345.10. $^1H$ NMR (500 MHz, $D_2O$) δ 8.91 (s, 1H), 7.53 (m, 4H), **7**.40 (s, 1H), 7.39 (s, 1H), 7.22 (m, 1H), 3.91 (s, 3H), 2.98 (m, 2H), and 1.81 (m, 2H).

**Preparation of (4-(((8-methoxyquinazolin-4-yl)amino)methyl)phenyl)phosphonic acid 56.**

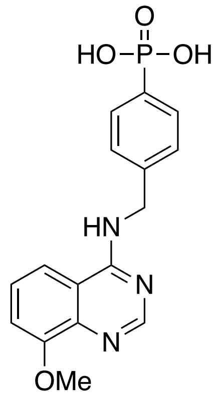

Prepared according to the same method as compound **23**. LCMS: $[M + H]^+$ *m/z* 346.10. $^1H$ NMR (400 MHz, $D_2O$) δ8.20 (s, 1H), 7.62–7.57 (m, 2H), 7.43–7.41 (m, 2H), 7.31–7.28 (m, 2H), 7.23–7.21 (m, 1H) and 2.93 (s, 3H).

**Preparation of (4-(((3-cyano-8-methoxyquinolin-4-yl)amino)methyl)phenyl)phosphonic acid 57.**

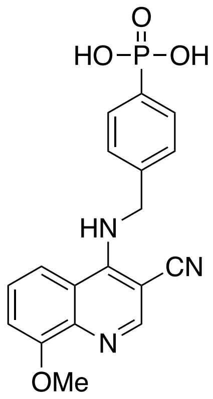

Prepared according to the same method as compound **23** to give **57** as an off-white solid. LCMS: $[M + H]^+$ *m/z* 370.10. $^1H$ NMR (400 MHz, $D_2O$) δ8.02 (s, 1H), 7.48–7.43 (m, 2H, 7.36–7.30 (m, 2H), 7.15–7.09 (m, 3H), 4.77 (s, 2H) and 3.81 (s, 3H).

**Preparation of (4-(((3-cyano-8-methoxyquinolin-4-yl)amino)methyl)benzyl)phosphonic acid 58**.

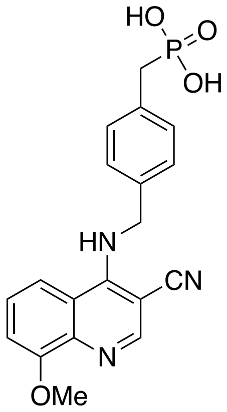

Prepared according to the same method as compound **23** to give **58** as a white solid. LCMS: $[M + H]^+$ *m/z* 384.15. $^1H$ NMR (400 MHz, Methanol-$d_4$) δ8.32 (s, 1H), 7.77 (d, $J$ = 8.4 Hz, 1H), 7.50 (t, $J$ = 8.4 Hz, 1H), 7.35–7.33 (m, 2H), 7.26 (d, $J$ = 7.6 Hz, 1H), 7.20 (d, $J$ = 8.4 Hz, 2H), 5.02 (s, 2H), 3.99 (s, 3H) and 2.85 (d, $J$ = 20 Hz, 2H).

**Preparation of (3-(1-(3-cyano-8-methoxyquinolin-4-yl)piperidin-4-yl)propyl)phosphonic acid 59.**

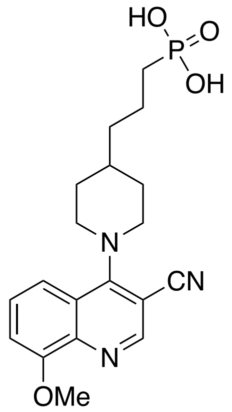


Prepared according to the same method as compound **14** to give **59** as a white solid. LCMS: [M + H]$^+$ *m/z* 390.20. $^1$H NMR (400 MHz, $D_2O$) δ8.30 (s, 1H), 7.39 (br s, 2H), 7.19 (br s, 1H), 3.98 (s, 3H), 3.73–3.70 (m, 2H), 3.30 (t, *J* = 12 Hz, 2H), 1.92–1.88 (m, 2H), 1.70–1.45 (m, 3H) and 1.40–1.27 (m, 6H).

**Preparation of (4-(((3-cyano-8-methoxyquinolin-4-yl)amino)methyl)phenyl)boronic acid 60.**

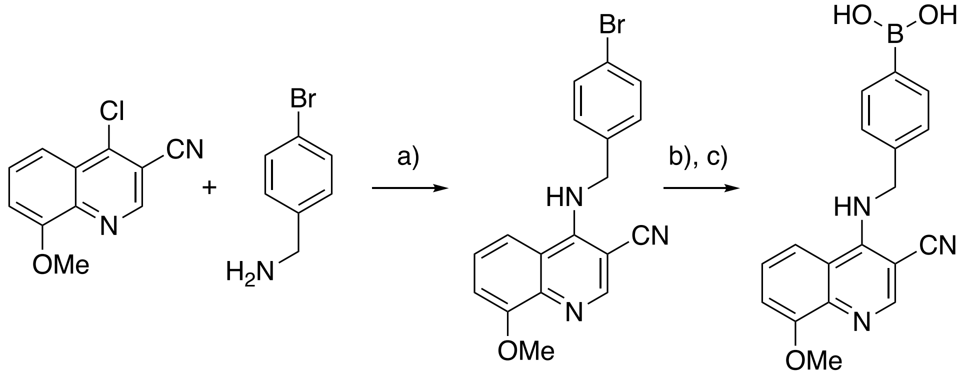


**105** **94** **106** **60**

a) $Et_3N$, $CH_3OCH_2CH_2OH$; b) $PdCl_2$(dppf), KOAc, $B_2Pin_2$, DMSO, 80 °C; c) HCl, EtOAc

To a solution of compound **105** (0.97 g, 5.0 mmol) in 2-methoxyethanol (10 mL) was added (4-bromophenyl)methanamine **94** (1.74 g, 10.0 mmol) and $Et_3N$ (1.51 g, 15 mmol). The mixture was heated at 100 ºC overnight, cooled to rt and then evaporated to dryness under reduced pressure. Chromatography (35% EtoAc in petroleum ether) gave **106** (1.5 g, 88%) as white solid. To a solution of compound **106** (69 mg, 0.2 mmol in DMSO (3 mL) was added bis(pinacolato)diboron (61.0 mg, 0.24 mmol), potassium acetate (58.8 mg, 3.0 mmol), Pd(dppf)$Cl_2$ (7.4 mg, 0.05 mmol). The reaction was degassed by purging with nitrogen and then heated at 80 ºC for 48 h. The mixture was cooled to rt, and diluted with ethyl acetate and then filtered through a pad of Celite®. The filtrate was evaporated to dryness under reduced pressure. The residue was dissolved in EtOAc (10 mL) and the HCl (4M, 0.2 mL, 4.0 mmol) solution in EtOAc was added. The mixture was stirred at rt overnight and then evaporated to dryness under reduced pressure. Chromatography [prep-HPLC (TFA)) gave **60** (40.5 mg, 65% over two steps) as white solid. LCMS: $[M + H]^+$ *m/z* 334.15. $^1$H NMR (400 MHz, Methanol-$d_4$) δ 8.69 (s, 1H), 7.90 (d, *J* = 8.4 Hz, 1H), 7.73 (t, *J* = 8.3 Hz, 2H), 7.61 (d, *J* = 8.1 Hz, 2H), 7.41 (dd, *J* = 17.3, 7.8 Hz, 2H), 5.05 (s, 2H), 4.13 (s, 3H).

**Preparation of (2-(1-(3-cyano-8-methoxyquinolin-4-yl)piperidin-4-yl)ethyl)boronic acid 61.**

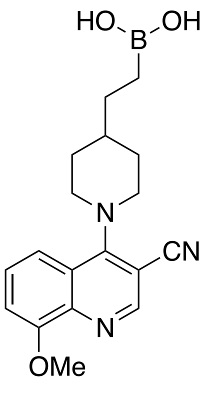

Prepared following the same procedure as compound **7**. Compound **61** was isolated as a yellow solid. LCMS: $[M + H]^+$ *m/z* 340.20. $^1H$ NMR (400 MHz, Methanol-$d_4$) δ8.81 (s, 1H), 7.83 (d, *J* = 12 Hz, 1H), 7.72 (t, *J* = 8 Hz, 1H), 7.60 (d, *J* = 8 Hz, 1H), 4.51 (d, *J* = 12 Hz, 2H), 3.81 (t, *J* = 12 Hz, 2H), 3.31 (s, 3H), 2.66 (s, 1H). 2.05 (br d, *J* = 12 Hz, 2H), 1.72–1.30 (m, 4H) and 0.91–0.85 (m, 2H).

**Preparation of 4-(((3-cyano-8-methoxyquinolin-4-yl)amino)methyl)-*N*-hydroxybenzamide 62.**

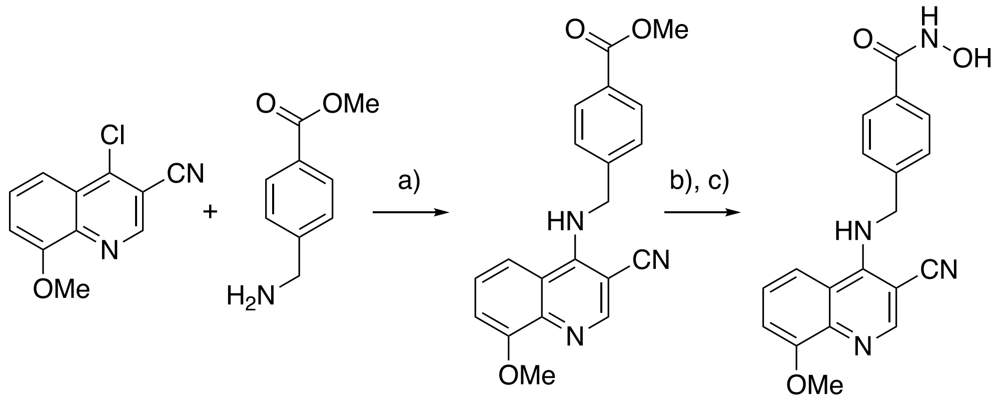


**105** **107** **108** **62**

a) $CH_3OCH_2CH_2OH$; b) NaOH, THF/$H_2O$; c) $NH_2OH$.HCl, BOP, DIPEA, THF

A solution of **105** (2.0 g, 8.9 mmol) and **107** (1.5 g 8.9 mmol) in 2-methoxyethanol (40 mL) was heated to reflux overnight and then cooled to rt. The reaction mixture was evaporated to dryness under reduced pressure and then triturated with EtOAc filtered and dried to give the crude compound **108** (1.7 g) as a light yellow solid.

To a solution of compound **108** (0.5 g, 1.55 mmol) in THF (20 mL) was added NaOH (0.17 g, 4.65 mmol, dissolved in 2 mL of water). The mixture was heated to 45 ºC overnight. The cooled solution was concentrated under reduced pressure and the residue treated with aqueous HCl (2N)

until pH 5.5 was realized. The resulting precipitate was filtered and dried to give the crude acid intermediate (0.3 g, 62% yield) as a light yellow solid. The crude acid was dissolved in DMF (10 mL) and then cooled to 0 ºC and placed under nitrogen. BOP (0.48 g, 1.06 mmol) and DIPEA (0.50 g, 3.88 mmol) were added followed by $HONH_2$-HCl (0.09 g, 1.26 mmol). The mixture was stirred at rt overnight, quenched with water (50 mL) and extracted with EtOAc. The organic phase was washed with water, and brine and dried ($Na_2SO_3$) and evaporated to dryness under reduced pressure. Chromatography (5% MeOH in $CH_2Cl_2$) and then Prep-HPLC ($H^+$, 0.1% TFA) gave **62** (34 mg, 10%) as off-white solid. LCMS: $[M + H]^+$ *m/z* 349.1 $^1H$ NMR (400 MHz, DMSO-*d6*) δ 11.19 (s, 1H), 9.06 (s, 1H), 8.55 (s, 1H), 8.45 (d, *J* = 8.7 Hz, 1H), 7.72 (d, *J* = 7.4 Hz, 2H), 7.39 (dd, *J* = 4.6, 2.9 Hz, 2H), 7.29 (dd, *J* = 12.3, 5.0 Hz, 2H), 5.11 (s, 2H), 3.93 (s, 3H).
